# Epigenetic inheritance of complex learning abilities in the mammalian brain

**DOI:** 10.1101/2025.08.31.673327

**Authors:** Samaa Zidan, Michael Andreyanov, Alaa Saleh, Noa Barnea, Sankhanava Kundu, Amit Kumar, Natanela Bakman, Tali Rosenberg, Maya Hovav, Friedrich Johenning, Dietmar Schmitz, Sadegh Nabavi, Steven Kushner, Asaf Marco, Shai Berlin, Jackie Schiller, Edi Barkai

**Affiliations:** Sagol Department of Neurobiology, Faculty of Natural Sciences, University of Haifa, Haifa, Israel; Department of Neuroscience, The Rappaport Faculty of Medicine and Research Institute, Technion-Israel Institute of Technology; Neuro-Epigenetics Laboratory, the Robert H. Smith Faculty of Agriculture, Food and Environment, the Hebrew University of Jerusalem, Israel; Neuroscience Research Center, Charité-Universitätsmedizin Berlin, Germany; Department of Molecular Biology and Genetics, Aarhus University, Denmark; Center for Proteins in Memory - PROMEMO, Danish National Research Foundation, Denmark; Department of Psychiatry, Columbia University, New York, NY 10032

## Abstract

For several decades, the question of whether cognitive and learning capacities can be inherited through non-genetic mechanisms has been the subject of ongoing debate. Here, we provide the first evidence of transgenerational inheritance of enhanced ability to learn complex tasks in the mammalian rodent brain. These inherited learning enhancements are not limited to specific stimuli, sensory modalities, or learning paradigms. Using behavioral, cellular biophysical, methylomics, genetics and molecular methods, we find that the inherited epigenetic modifications reflect an enhanced neuronal learning state, driven by increased intrinsic neuronal excitability in most pyramidal neurons in the relevant neuronal networks.

This enhancement is mediated by persistent downregulation of the muscarinic M-current and is associated with widespread changes in DNA methylation, notably within coding genes associated with the M-current, the Kv7 pathway, in the hippocampi of trained F0 rats, as well as in non-coding RNAs in their sperm samples. Remarkably, a significant portion of these DNA methylation changes were also observed in the hippocampi of their untrained F1 offspring. These findings suggest that complex learning abilities can be inherited in the mammalian brain, as the offspring of trained rodents are born with the biophysical modifications that enable them to become super-learners, the exact change that occurs in their parents’ brains only after the rule learning.

## INTRODUCTION

It is accepted that the influence of parents over their offspring’s cognitive abilities, learning capacities, and rule-formulation, to name a few, are mediated through two primary mechanisms, namely hereditary-(nature) and environmental (nurture) factors. Whereas the former involves the transfer of genetic material from parents (ancestral generation, F0) to offspring (F1) to directly define the content of the offspring’s genome, the latter describes non-genetic and non-heritable features (e.g., cultural and socio-economical) that are known to have profound influences on the behavior of the offspring. Over the past two decades, a third mechanism has gained recognition—the inheritance of non-canonical genetic factors that can substantially influence or shape offspring behavior. Accumulating evidence indicates that experience-dependent traits may be transmitted across generations specifically via epigenetic modifications in the parental genome, passed through the gametes to the offspring^1–9^.

Epigenetic transmission of acquired behavioral traits has been extensively demonstrated in simpler organisms, notably *Caenorhabditis elegans* and *Aplysia*. In these species, inherited behavioral modifications include sexually dimorphic traits linked to mating success, as seen in *C. elegans*^10,11^, responses to ancestral starvation, which enhance survival under nutrient-scarce conditions^10^, and memory retention of simple training paradigms in *Aplysia*, where learned sensitization and habituation are transferred across generations^12,13^. These findings offer compelling evidence that environmental experiences can modify the parental genome in ways that are heritably transmitted to the next generation, influencing behavior through non-genetic mechanisms of inheritance.

In mammals, however, epigenetic inheritance has primarily been observed in pathological and extreme conditions. These include transgenerational effects of early-life trauma and chronic stress^2,7,14^, metabolic disorders influenced by parental diet and health^15^, increased disease susceptibility through inherited epigenetic modifications^16^, and even fear-conditioning responses to specific odors which may be passed onwards through sperm-mediated epigenetic modifications^17,18^. Importantly, whether epigenetic inheritance can also extend to more generalized learning abilities remain unknown. Given the emerging role of neuronal epigenome remodeling as a critical determinant of learning capacity^1^, alongside evidence that learning and memory processes can drive persistent epigenetic modifications within neuronal populations ^12,13,19^, it is plausible that complex physiological traits—including complex learning abilities—could be heritable. This hypothesis suggests that experiences leading to optimized neural plasticity and cognitive function may be epigenetically transgenerationally inherited.

In this study, we investigate whether complex learning abilities—specifically, the capacity for generalization in rule-learning—can be transmitted across generations. Furthermore, we explore whether this inheritance is mediated by an epigenetic mechanism. Given that defined neuronal cellular-biophysical changes have been previously identified as critical to rule-learning^20–29^, we assessed whether these modifications are transgenerationally inherited, thereby endowing offspring of trained parents with cognitive abilities that are not typically observed in naïve animals.

Leveraging a multidisciplinary approach that combines behavioral analysis, methylomics, genetics, and biophysics, we provide evidence that exceptional learning capabilities, acquired by rats and mice in the parental (F0) generation through intensive rule-learning, are epigenetically transmitted to naïve offspring across multiple generations (F1-F3). Briefly, we employed the well-established olfactory discrimination (OD) task^20–25^. Training rodents in this particularly demanding task triggers a phase of accelerated learning for additional odors, to which the animals were not exposed during the training phase. This is evidenced by a significant enhancement in their ability to rapidly acquire and retain new odor memories after mastering the initial task. Such a phenomenon strongly suggests that the animals have undergone rule learning, enabling them to generalize acquired learning strategies to novel stimuli^22–25^. Strikingly, animals that were trained in the complex OD task also show enhanced learning capability in the Morris water maze^26^. Thus, mastering ‘rule-learning’ leads to a broad improvement in overall learning ability. We further find that rule-learning is accompanied by an increase in intrinsic neuronal excitability of pyramidal neurons, mediated by persistent reduction in the medium and late after hyperpolarization (AHPs), consistent with previous reports in various brain regions (piriform cortex^21,27^, hippocampus^26^ and amygdala^28^). The sustained reduction of the AHP necessitates continuous activation of PKC and ERK^29^, ultimately leading to the downregulation of the cholinergic M-current^27^. Of note, AHP reduction and the resulting increase in neuronal excitability do not serve as the direct mechanism for encoding specific sensory memories or event sequences. Instead, they establish a neural state described as a “learning mode,” in which neuronal ensembles become primed for enhanced plasticity. The behavioral outcome of this state is a significant improvement in learning performance, specifically in tasks that critically depend on these neuronal networks^20,26^.

Our findings challenge conventional boundaries of transgenerational inheritance of learned ability and our model system fundamentally advances previous research on transgenerational inheritance of learned behaviors in four key manners. *First*, prior studies primarily focused on the transmission of simple learned behaviors, such as conditioned fear or aversive avoidance responses, whereas our paradigm investigates the heritability of generalized cognitive ability. *Second*, most transgenerational learning studies report the inheritance of responses to specific stimuli (e.g., odor-associated fear conditioning), whereas we demonstrate the transmission of a generalized learning capacity. *Third*, inherited learning ability correlates with increased intrinsic neuronal excitability, mediated by persistent downregulation of the cholinergic-muscarinic potassium current (M-current). *Fourth*, we identify epigenetic modifications within the muscarinic gene pathway, specifically in the Kv7 pathway, marked by altered DNA methylation patterns in both trained F0 animals and their naïve F1 progeny. This aligns with emerging evidence that neuronal excitability serves as a fundamental determinant of cognitive plasticity, with epigenetic mechanisms playing a crucial role in sustaining learned adaptations across generations.

## RESULTS

### Transgenerational inheritance of learned ability

We trained male and female rats from the ancestral (F0) generation using two distinct behavioral paradigms: the complex odor discrimination (OD) task and the Morris water maze (WM) (**Fig. 1a**)^30^. Training continued until F0 animals successfully acquired rule-learning (F0-OD or F0-WM) (see **Methods**). Animals from the F0 generation reached the criterion of OD rule in 7.4±0.29 days (n= 12 rats), consistent with previous reports^20,21^. Trained F0 animals were then bred with other trained F0 animals (from same task) and their F1-F3 offspring were assessed for learning capacity across various tasks. Offspring performance was compared to that of progeny from either untrained (naïve) or *pseudo*-trained animals, namely animals that underwent identical training but received rewards at random (**Fig. 1b**). Notably, F1–F3 animals did not receive any training prior to assessment of their learning abilities. We observed that non-trained F1 offspring (briefly, F1 offspring) of OD-*trained* parents exhibited significantly improved success rate for odor discrimination (**Fig. 1b, red closed circles**), compared to age-matched F1offspring of naïve- or *pseudo*-trained rats (**Fig. 1b, blue and black circles, respectively**). The F1 progeny of OD-trained parental rats achieved criterion performance in under four days, whereas the offspring of naïve or *pseudo*-trained parents required approximately twice that duration to reach the same level of task proficiency (**Fig. 1c**). The observation that the F1 progeny of *pseudo*-trained animals performed similarly to those of the naïve group suggests that the transgenerational enhancement in learning observed in the F1 progeny of trained animals cannot be attributed to mere exposure to the maze environment, odor cues, or handling. Rather, it suggests the presence of a specific transgenerational mechanism^26^ driven by the successful acquisition of task rules and genuine learning in the parental generation. This hypothesis was strengthened by the observation that the offspring of WM-*trained* animals outperformed the other groups in the OD-task; a task their parents had never encountered (**Fig. 1b, orange open circles**). The inverse was also found to be true, namely F1-offspring of F0 OD-*trained* parents showed faster acquisition of the WM task compared to F1 offspring from naïve- or *pseudo*-trained F0 parents (**Fig. 1d**). Remarkably, transgenerational mechanism of learning ability persisted in the F2 and F3 generations, in which case only the F0 generation underwent training (or *pseudo*-training) on the OD task (**Fig. 1e**).

**Figure 1.**
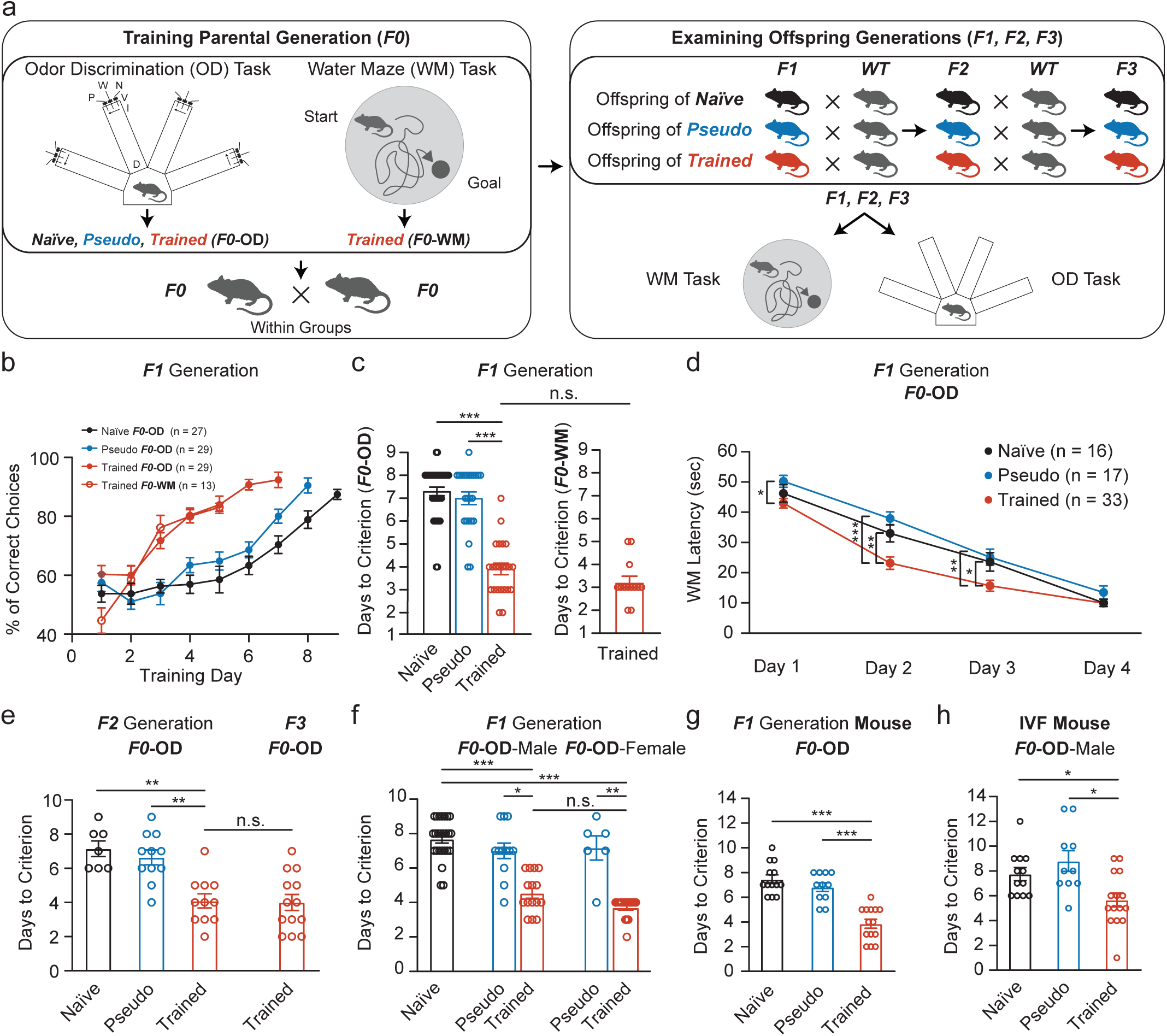
Enhanced learning by offspring of trained rats and mice. **(a)** Cartoon depiction of the behavioral paradigms. (**Left**) The ancestral generation (F0) of rats (or mice, below) undergoing training (Trained group) in the odor discrimination task (OD) or Morris water maze task (WM), compared to Naïve or Pseudo-trained (pseudo) animals. The F0 animals were then bred within groups. (**Right**) Offspring (F1) of Fo-Naïve, F0-Pseudo and F0-Trained animals were further bred to F2 and F3 generations with wildtype (WT) animals or were examined by either the OD task or the WM task. Note-F1-F3 did not undergo any training. **(b)** Learning curves of F1 offspring from F0 Naïve-, Pseudo- or Trained-rats in the odor discrimination task (black, cyan and orange closed circles, respectively). F1 offspring from trained parents (orange, closed circles) reach maximal performance (80% correct choices) twice faster than the other groups; summarized in (**c, left**). Enhanced learning capability was also apparent when the parental generation was trained in the WM task (orange, open circles); summarized in (**c, right**). **(d)** Offspring of OD-trained rats show enhanced learning capabilities in the WM task. **(e)** Enhanced learning capabilities is persistent in the F2 and F3 offspring from F0-Trained rats. **(f)** Enhanced cognitive performance is observed in F1 progeny following the training of a single parent—either male or female rat. **(g)** Transgenerational inheritance of learning ability extends to mice. **(h)** Offspring of Naïve mothers fertilized with sperm from OD trained F0 males show enhanced learning capability. Data are presented as mean ± SEM. *, P<0.05; **, P<0.01; ***, p<0.001; n.s., non-significant.

To determine whether the transgenerational effect was influenced by the parental sex, we selectively trained either male or female F0 animals and bred them with naïve F0 partners. F1 offspring of trained males or females learned the OD task equally as fast, but significantly faster than those of naïve or *pseudo*-trained F0-males (**Fig. 1f**). Lastly, and complementary, we cross-fostered F1-progenies from trained F0 mothers with *pseudo*-trained mothers (and vice versa) to explore whether maternal behavior would have any effect over the phenomenon, but noted that fostering had no effect and that the F1-offpsring remained the best performers regardless having been raised by *pseudo*-trained mothers (**Fig. 1f**). These demonstrations suggest that transgenerational inheritance of learning ability occurs through both the male and female germline (see below) and further excludes maternal behavior and other social cues as the source of the effect, as previously shown for other behavioral traits^31^.

### Transgenerational inheritance of learning ability extends to mice

We next investigated whether the transgenerational inheritance of learning ability would extend to another mammalian species—specifically, mice. While rats and mice share many physiological and anatomical features, it is increasingly recognized that rats are not simply “larger mice”^32^. The two rodent species diverged approximately 12 million years ago^33^, and this evolutionary separation is reflected in numerous neurobiological distinctions. For instance, differential expression of genes (e.g., 4,700 genes are differentially expressed in hippocampal neurons)^32,34^, cell types in brain regions^35^, and distribution of neurotransmitter receptor subtypes across the brain^36^. Such molecular and anatomical differences manifest in distinct behavioral and cognitive profiles, making them highly suitable for assessing generalization of cognitive phenomenon.

We therefore trained the parental F0 generation of mice and find that their F1 offspring required significantly fewer days to reach the learning criterion compared to the control groups, as observed in rats (**Fig. 1g**). This positive finding in mice thereby enabled us to then directly explore whether the mechanism would be transmitted through the germline (and to exclude other post-conception, experience-dependent plasticity mechanisms) by *in vitro* fertilization (IVF) (IVF-derived embryos are particularly challenging in rats^37^). We produced F1 offspring via using sperm from OD-*trained*, *pseudo*-trained or naïve F0 male mice (collected 10 days after the trained mice reached the learning criteria). Oocytes for IVF were obtained from superovulated naïve F0 female mice and implanted into an independent group of naïve F0 female mice (**methods**). We find that F1 offspring conceived exclusively from the sperm of OD-trained F0 males reached the learning criterion significantly faster compared to F1 offspring derived from the sperm of naïve or *pseudo*-trained F0 males (**Fig. 1h**). These results demonstrate that enhanced learning ability observed in the offspring of trained animals is not species-specific and confirm that the effect is transmitted through the sperm of trained males, supporting our above observation regarding transmittance via the male germline in rats (see **Fig. 1f**). Furthermore, these results particularly rule-out maternal influences or post-conception experiences as confounding factors. These combined observations strongly support the existence of a germline-dependent, experience-induced mechanism of inheritance of cognitive traits.

### Higher neuronal intrinsic excitability is passed on to F1-F3 generations from parents’ brains who acquired rule leaning

To investigate the biophysical mechanism underlying the transgenerational inheritance of learned ability, we chose to focus on post-burst afterhyperpolarization (AHP)-mediated changes in intrinsic excitability. First, this process is consistently implicated in the cellular encoding of learning and memory across multiple brain regions, including the hippocampus, piriform cortex, and amygdala^20,21,26,2^. For instance, a reduction in AHP following learning enhances firing responsiveness of excitatory neurons, facilitating synaptic integration and consolidation of memory traces^20,21,26,28^. Second, this form of learning-induced intrinsic plasticity is both experience-dependent and long-lasting, making it a plausible candidate for mediating durable, intergenerational changes in cognitive function. In contrast to synaptic mechanisms such as long-term potentiation (LTP), which are often circuit-specific and require external stimulation paradigms, changes in intrinsic excitability are cell-autonomous, more broadly distributed, and potentially more amenable to germline-mediated modulation via epigenetic mechanisms (e.g., ^1^). Given these characteristics, we reasoned that examining intrinsic excitability through AHP modulation would provide a mechanistically tractable and biologically relevant entry point for understanding how learning experiences in the parental generation could become encoded into transmissible physiological states in their offspring.

*Ex vivo* recordings from rat hippocampal CA1 pyramidal neurons from non-trained F1 offspring of OD-trained, naïve and *pseudo*-trained F0 parents showed that the medium and late AHPs were significantly reduced in offspring of OD-*trained* parents, compared to offspring of naïve or OD pseudo-trained parents (**Fig. 2a-c**). No differences were observed in resting membrane potential or input resistance between the groups (**Suppl. Fig. 1**). Similar reductions in AHPs were maintained in hippocampal CA1 pyramidal neurons from F2 and F3 offspring of OD-*trained* F0 ancestors exclusively (**Fig. 2d**). We further find a strong and negative linear relationship between medium and late AHP’s amplitudes, and the number of action potentials generated in response to a fixed somatic current injection in CA1 pyramidal neurons (**Fig. 2e, f**), consistent with the key role of medium and late AHPs in suppressing repetitive spike firing^27,38,39^. Indeed, the average number of action potentials evoked in neurons from trained rats’ offspring in a sequence of 17 stimuli was significantly higher than in neurons from naïve and pseudo-trained offspring (**Fig. 2g, h**).

**Figure 2.**
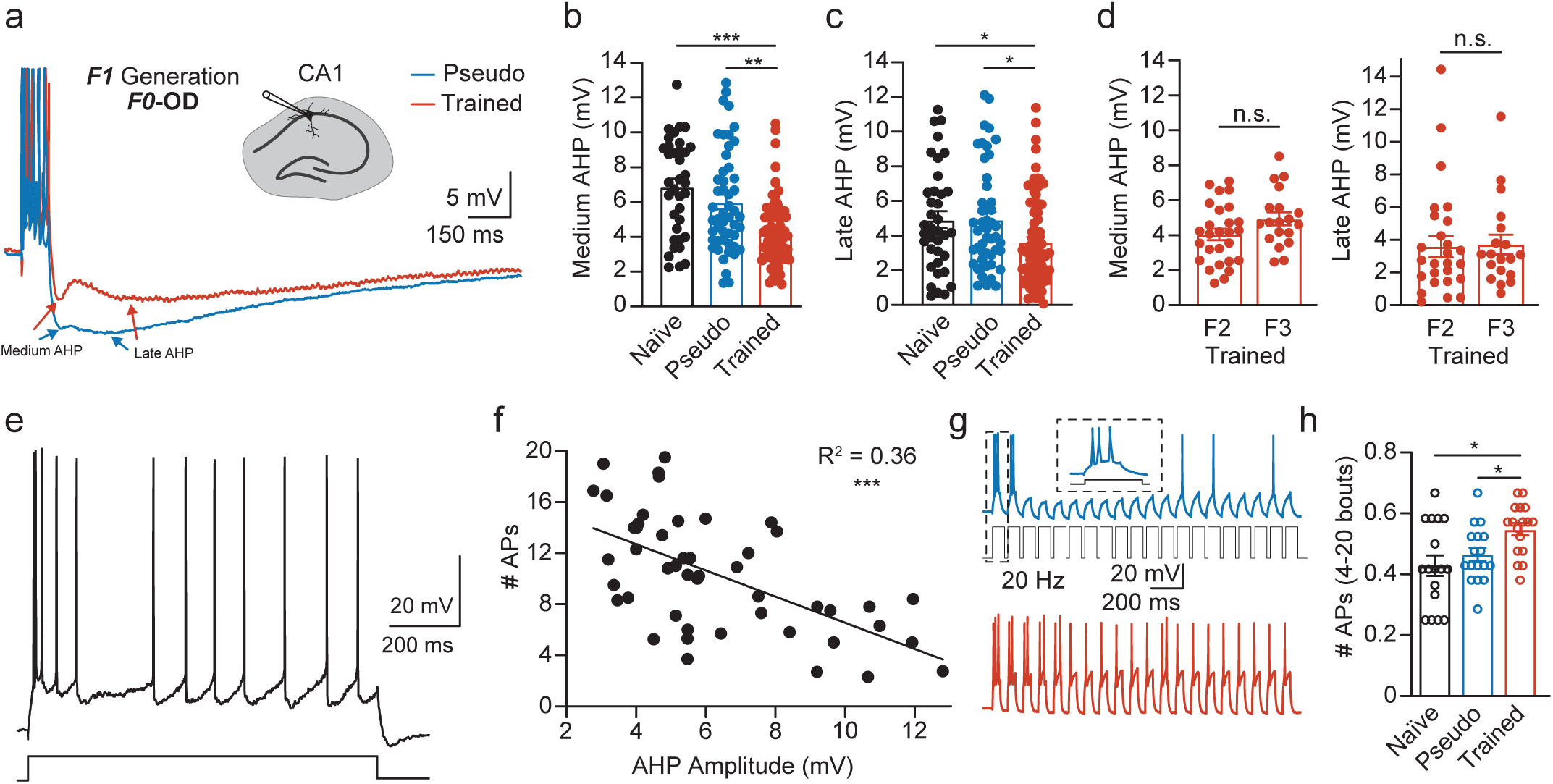
Enhanced intrinsic excitability in CA1 pyramidal neurons from the F1-progeny of F0-Trained rats. **(a)** Representative traces showing post burst AHPs recorded in neuron from progenies of F0-pseudo-(cyan) and F0-Trained (orange) parents. Arrows indicate medium and late AHPs. *Inset:* location of the recording electrodes. **(b, c)** The amplitudes of the medium and late AHPs are significantly reduced in F1 progeny derived from trained animals (black), compared to pseudo-trained and naïve groups. **(d)** The amplitudes of the medium and late AHPs are maintained repressed in neurons from F2 and F3 descendants of trained rats. Dots note the AHP value in each neuron. **(e)** Representative trace showing neuronal firing to current injection (1 sec, at intensity-I_th_x2; 1.25 nA in this neuron). **(f)** Negative association between AHP amplitude and number of action potentials (# APs). Results are shown for 48 neurons. A highly significant negative correlation between the AHP amplitude and the number of action potentials is apparent (n= 48 neurons; r= -0.6). **(g)** Representative traces from CA1 neurons recorded from neurons of progenies from F0-pseudo (cyan) or trained animals (orange), in response to 20 repetitive stimulating bouts (see **methods**). Neurons from the progenies of the F0-pseudo-trained rats mostly show firing at the onset of the protocol (at the first two bouts), and typically fail to generate repetitive spiking at subsequent bouts, whereas neurons from trained parents exhibit firing throughout the protocol, summarized in **(h)**. **(h)** Average of the number of action potentials generated by neurons in stimulation bouts 4-20. Data are presented as mean ± SEM. *, P<0.05; **, P<0.01; ***, p<0.001; n.s., non-significant.

### Neurons in F1-offspring of trained fathers are more excitable due to persistent reduction of the M-current

Olfactory rule learning promotes an enhancement of intrinsic neuronal excitability by reduction of the cholinergic-dependent KCNQ current (the muscarinic (M)-current driven by KCNQ)^40–43^, leading to reductions in medium and late post-burst AHPs (**Suppl. Fig. 2**)^20–22,44–46^. However, long-lasting attenuations of AHP are maintained by persistent PKC-dependent inhibition of KCNQ^21,27,29^. Notably, though well established in cells of the trained animals themselves, this phenomenon has not been explored in offspring of trained animals. To investigate whether the M-current pathway is engaged in F1 offspring of trained animals, we first probed the PKC-dependent inhibition of KCNQ current. We inhibited PKC activity via the cell-permeable GF109203X, which had no effect on AHP amplitude in cells from offspring of naïve and pseudo-trained rats (**Fig. 3a-c**). In contrast, inhibition of PKC restored the medium- and late AHP in CA1 neurons from F1 offspring of *trained* rats to a comparable level as for F1 offspring of naïve or pseudo-trained rats (**Fig. 3b, c**). These data suggest that the long-lasting medium and late AHP reductions in neurons of F1 offspring from trained parents are mediated by persistent PKC-dependent inhibition of the M-current.

**Figure 3.**
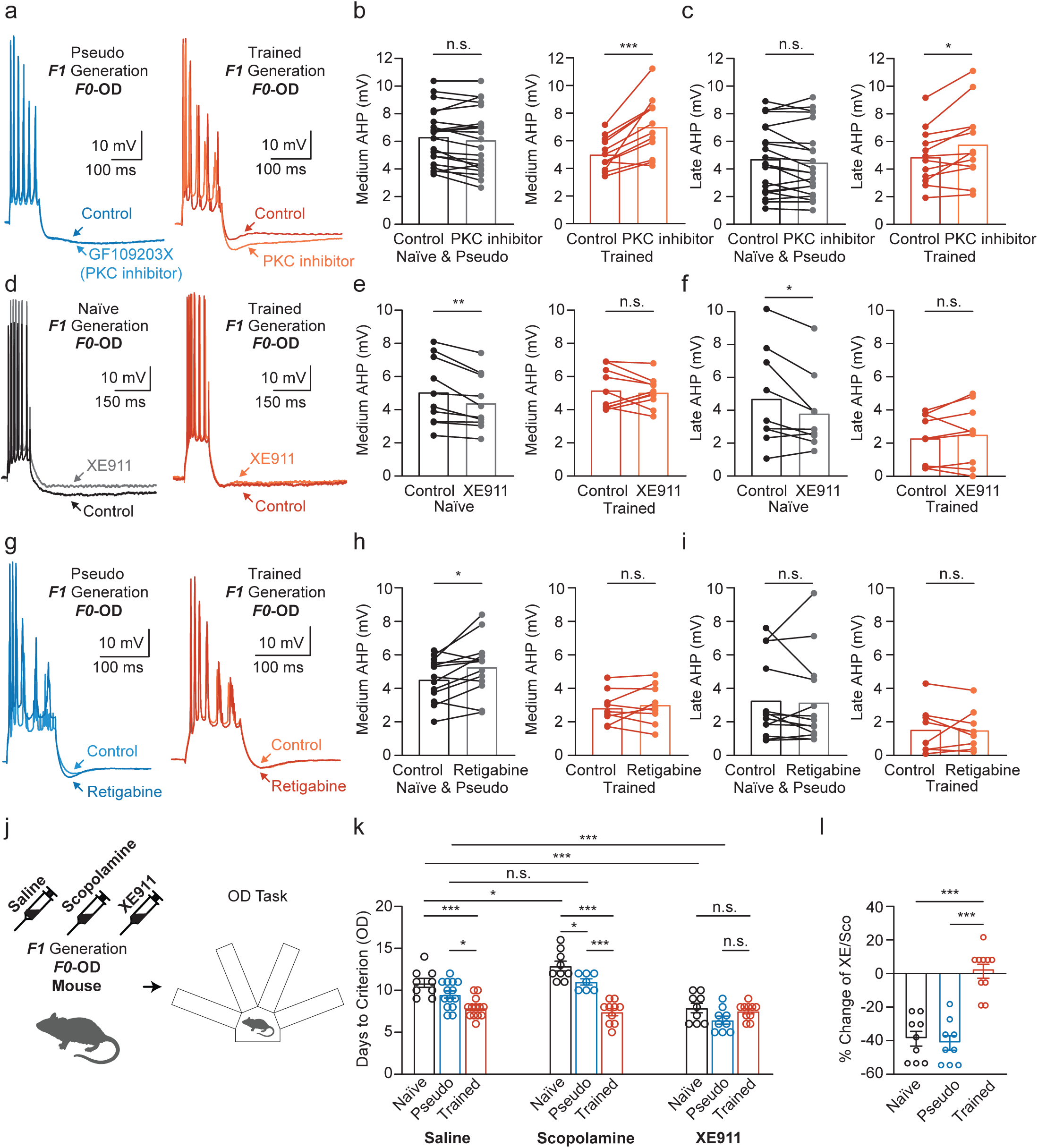
Inherited enhanced neuronal excitability is mediated by persistent PKC-dependent down regulation of M-current. **(a)** Representative traces from neurons from progenies of F0-Pseudo (cyan and blue) or F0-trained parents (orange and red), before and after application of GF109203X (a widely used PKC inhibitor). Inhibition of PKC has no effect over the amplitudes of the medium and late AHP in F1 from naïve and pseudo-trained animals but produces a strong increase in the amplitudes of the medium and late AHP in the F1 progenies from Trained animals, summarized in **(b)** and **(c)**, respectively. **(d)** Representative traces from neurons from progenies of F0-Naïve (black and grey) or F0-trained parents (orange and red), before and after application of XE911 (a selective blocker of KCNQ, Kv7). Blocking of KCNQ reduces the amplitudes of the medium and late AHP in F1 from Naïve-trained animals but shows no effect over the amplitudes of the medium and late AHP in the F1 progenies from Trained animals, summarized in **(b)** and **(c)**, respectively. **(g)** Representative traces from neurons from progenies of F0-Pseudo (cyan and blue) or F0-Trained parents (orange and red), before and after application of retigabine (a selective channel opener of KCNQ, Kv7). Activation of KCNQ increases the amplitude of the medium, but not late AHP in F1 from Pseudo-trained animals and has no effect over the amplitudes of the medium and late AHP in the F1 progenies from Trained animals, summarized in **(h)** and **(i)**, respectively. **(j-l)** Modulation of cholinergic activity *in vivo* in mice. **(j)** Schematic representation of F1 offspring of Trained parents administered with saline, scopolamine (muscarinic receptor antagonist) or XE991 and probed on the OD task. **(k)** F1 progenies of Trained-parents are indifferent to manipulations of the cholinergic M-current pathway (orange bars). Conversely, Scopolamine decreases performance of F1 progenies from Naïve parents with no effect over offspring of Pseudo-trained parents. XE991 improves the performance of Naïve and Pseudo groups—without affecting the Trained group—bringing their performance to comparable levels to those observed in F1 progeny of F0-Trained parents. **(l)** Summary of the effect of the M-current blocker compared to inhibition of the muscarinic receptor, showing that F1 progenies from Naïve and Pseudo-trained mice require ∼40% less time to complete the task, compared to mice treated with the inhibitor of the muscarinic receptor. In contrast, the two treatments had no effect on F1 progenies from the F0-Trained group. Data are presented as mean ± SEM. *, P<0.05; **, P<0.01; ***, p<0.001; n.s., non-significant.

These motivated us to directly explore the involvement of channel activity in the phenomenon. To this end, we applied a selective KCNQ blocker (XE991, 10 µM), which expectedly decreased the medium and late AHPs’ amplitudes in neurons from control naïve offspring. However, this treatment had no effect on neurons from F1 offspring from the trained parents (**Fig. 3d-f**). Analogously, we applied a KCNQ channel activator (retigabine, 10 µM) that significantly enhanced the medium AHP in neurons from naïve offspring, whereas had no effect over neurons from the F1 offspring of trained animals (**Fig. 3g-i**). This treatment, however, had no effect on the late AHPs in both groups (**Fig. 3i**). Both medium and late AHP were resistant to the activation in F1 offspring from trained animals. It is worth mentioning that the lack of increase in late AHP by retigabine, in neurons from offspring of naïve and pseudo-trained rats, contrasts the results obtained with application of the PKC blocker that did influence both types of AHP (compare **Fig. 3a and h**). These suggest that late AHP requires PKC activity and cannot be simply imitated by a channel opener. This is consistent with reports showing that long lasting attenuations of AHP result from persistent PKC-dependent inhibition of KCNQ^21,27,29^. Thus, despite the presence of the channel in neurons of F1 offspring from trained animals, the channel is suggested to be persistently inhibited by PKC (**Fig. 3a, red trace**). Furthermore, the lack of enhancement of the late AHP in control cells following direct activation of the channel (**Fig. 3h, I; naïve, black**), suggests a secondary mechanism that converges onto PKC activation. These strongly complements the above observations with the M-channel blocker (**Fig. 3d-f)**, in which instance AHP could not be further attenuated, suggesting the channel is already intrinsically inhibited in neurons of F1 offspring from trained parents.

These findings confirm that in F1 offspring of trained animals, the M-current is persistently suppressed via a PKC-dependent mechanism. Importantly, the fact that the PKC blocker could restore both AHP components, whereas the channel opener only affected the medium AHP in control neurons, implies that additional upstream pathways may be involved in activating and maintaining this PKC-dependent suppression. This prompted us to explore alternative signaling mechanisms that could converge onto PKC and contribute to the persistent suppression of the M-current. One such candidate is the GluK2 kainate receptor pathway. In several brain regions, GluK2 activation has been shown to reduce AHP and enhance intrinsic excitability via PKC- and Ca²⁺-dependent signaling cascades. Given that olfactory learning enhances glutamatergic transmission, and that kainate receptors are strongly expressed in hippocampal circuits, it is plausible that GluK2 activation may contribute to the transgenerational inheritance of AHP modulation. Indeed, activation of GluK2 has been demonstrated to be both necessary and sufficient for OD rule learning by suppressing AHP via modulation of K⁺-channels through the PKC/MAPK cascades^25,47^. Thus, we next examined whether GluK2 signaling contributes to the persistent PKC-mediated inhibition of the M-current observed in the offspring of trained animals. We indeed find reduced AHP is maintained in F1-offspring of trained parents via GluK2 activation as in the parents’ brain after rule learning (**Suppl. Fig. 3a-c**). Also, synaptic stimulation (leading to GluK2 activation) recapitulates the observations with kainate application, showing that the AHP in neurons from F1-offspring of trained parents is inherently and robustly attenuated (i.e., cannot be further reduced)^29,48,49^ (**Suppl. Fig. 3d-f**). Given the commonality of synaptic- and GluK2 activation, and AHP modulation, it was reasonable to propose that these mechanisms would not limited to the neuronal CA1 pyramidal cell population^28^. We thereby examined the medium AHP in three additional brain regions: hippocampal CA3, basolateral amygdala (BLA) and layer II neurons of the medial prefrontal cortex (mPFC) (**Suppl. Figs 4 and 5**). In all cases, we find strong attenuation of the medium AHP and further show that the late AHP amplitude was particularly and strongly attenuated in CA3 neurons from trained rats’ offspring. Notably, transgenerational inheritance of generalized learning ability extends to additional mPFC- and BLA-dependent tasks (**Suppl. Fig. 5**).

### Modulation of cholinergic activity affects rule learning in F1 offspring of trained mice

To determine whether cholinergic suppression of the M-current contributes to the transgenerational inheritance of enhanced learning, we pharmacologically manipulated this pathway *in vivo*. F1 offspring of trained mice were systemically administered either saline, the muscarinic receptor antagonist scopolamine, or the M-current blocker XE991 (**Fig. 3j**). As expected, offspring of trained animals learned the task more rapidly than those from naïve or pseudo-trained parents, regardless of saline or scopolamine treatment (**Fig. 3k**). Interestingly, scopolamine impaired learning performance in F1 offspring of naïve mice, and to a lesser extent in pseudo-trained offspring, when compared with saline treatment. These suggest that muscarinic signaling is important for baseline learning in untrained lineages. Moreover, the performance of scopolamine-treated pseudo-trained offspring was significantly different from that of scopolamine-treated naïve offspring, indicating subtle training-related changes even in the absence of learning success in the parents. Overall, the results demonstrate that inhibition of the cholinergic pathway increased the time required to achieve rule learning in F1 naïve animals, while having no effect on trained animals— highlighting that cholinergic modulation via the M-current plays a selective role in learning capacity that can be modified by parental experience and potentially transmitted across generations.

Unexpectedly, blocking the M-current with XE991 eliminated the performance differences between offspring groups. F1 offspring of trained, naïve, and pseudo-trained mice all learned the olfactory discrimination task in a similar timeframe (**Fig. 3k, right**). XE991 significantly enhanced learning in naïve and pseudo-trained offspring, reducing their time to criterion compared to scopolamine controls. In contrast, XE991 had no effect on trained offspring, indicating their enhanced learning is independent of M-current modulation (**Fig. 3l**). Thus, persistent suppression of the M-current can elevate learning in untrained lineages, supporting its role in the transgenerational inheritance of enhanced cognitive ability.

### Transgenerational inheritance of learning ability is associated with distinct patterns of hippocampal DNA methylation

Our behavioral and mechanistic data suggest that enhanced learning ability can be transmitted across generations following parental training. Given that no genomic sequence changes are expected from training alone, epigenetic regulation offers the most plausible mechanism. Indeed, DNA methylation is well established as capable of shaping offspring behavior by altering gene expression patterns (behavioral-inheritance)^1,4,5^, such as epigenetic inheritance of susceptibility to disease and pathologies following parental exposure to stressors^3,6,7^. To examine whether the transgenerational transmission of enhanced learning involves experience-dependent DNA methylation changes passed through the germline, we conducted Reduced Representation Bisulfite Sequencing (RRBS) on hippocampal and sperm samples from trained, pseudo-trained, and naïve F0 fathers, as well as from their naïve F1 offspring. Briefly, methylated sites, meeting quality control criteria across all samples were subjected to paired differential methylation analysis in F0 and F1 (Naïve vs. Trained, Naïve vs. Pseudo and Trained vs. Pseudo, percentage change in methylations > %2.5, FDR<0.01). Given their substantial impact on gene transcription regulation, we opted to focus our analysis on differentially methylated regions (DMRs); contiguous genomic segments (of 1000 bp) demonstrating consistent variations (see **Methods**).

Our genome-wide methylation analysis revealed that OD training in F0 animals induced widespread epigenetic remodeling with a bias toward hypomethylation compared to naïve controls (1387 hypo- vs. 1159 hyper-DMRs) (**Fig. 4a**) or pseudo-trained animals, (5309 hypo- vs. 675 hyper-DMRs) (**Fig. 4b**), whereas pseudo-training alone instigated a striking predominance of hypermethylation (4944 hyper- vs. 731 hypo-DMRs) (**Fig. 4c**). This pattern suggests that rule learning—unlike passive exploration, exposure or handling, drives a distinct epigenetic trajectory marked by targeted demethylation, aligning with growing evidence that behavioral plasticity is encoded through selective demethylation of learning-relevant genomic regions^50,51^.

**Figure 4.**
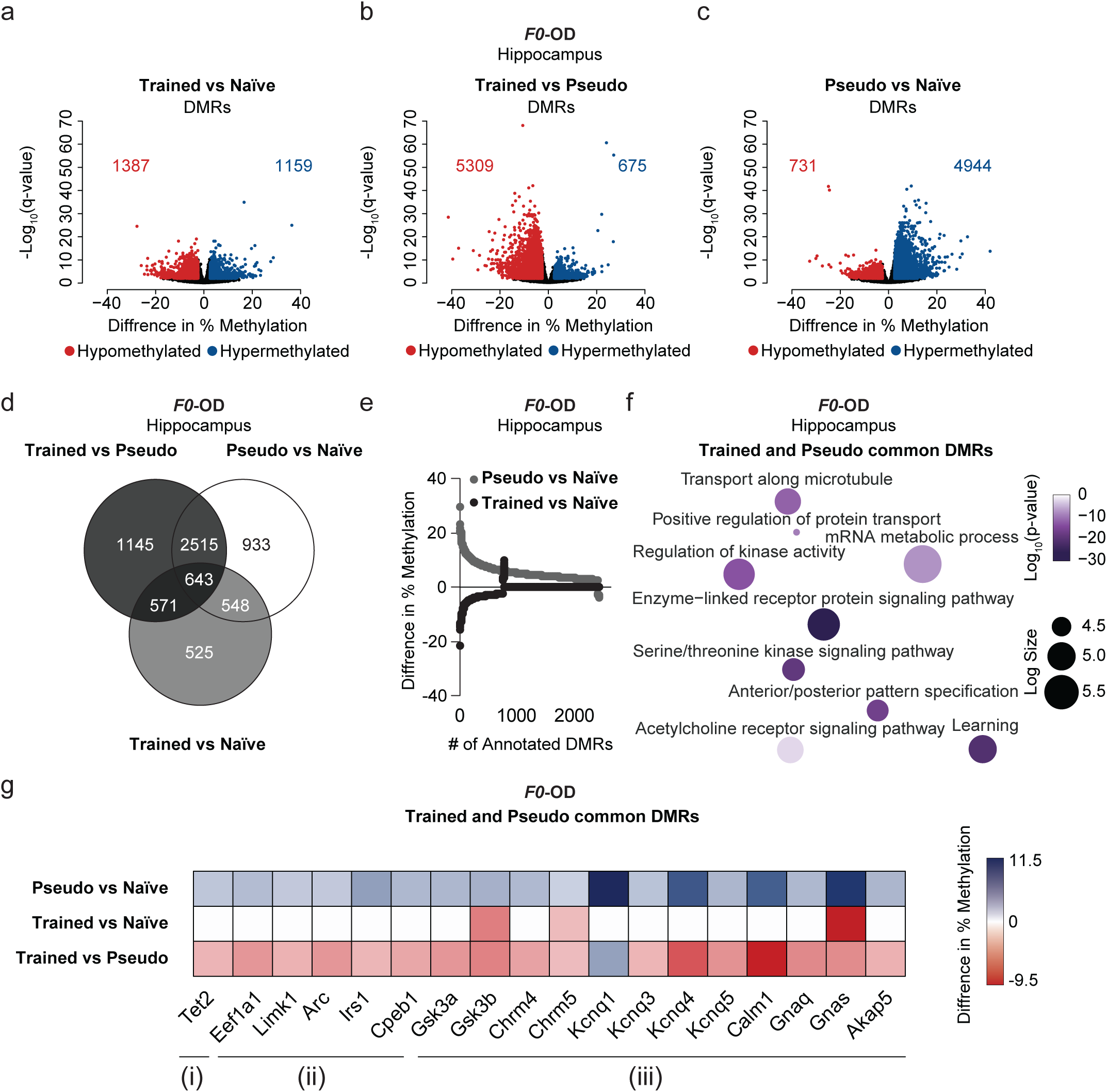
Superb learning capabilities in F0 trained rats is associated with hippocampal hypomethylation of genes linked to modulation of PKC-dependent pathways and the M-current. **(a-c)** Volcano plots of hippocampal differentially methylated regions (DMRs). Values of methylation from pairwise comparisons (difference in % methylation) were plotted against the average log10(*p-value*). Differentially hypo-methylated DMRs are shown in red, hyper-methylated DMRs in blue, and non-significantly methylated DMRs in black (and in color-coded numbers). Cutoff criteria – percentage change in methylations > +/-2.5, FDR<0.01, *n* = 2 /per group. **(d)** Overlap of DNA methylation profiles represent common or unique signatures between pseudo and trained animals. Venn diagrams (via InteractiVenn; see **methods**) was shows overlaps between annotated genes to DMRs. **(e)** common -DMRs between F0-Pseudo and -Trained rats. Each dot represents percentage of change in DMR levels from the pair-wise comparison to naïve group. **(f)** Gene ontology (GO) analysis of common DMRs (via Metascape and Revigo; see **methods**). Color indicates *p-value* and circle size indicates log enrichment size scale. **(g)** Heatmap showing representative genes annotated to DMRs extracted from the common-DMR GO analysis. Rows represent percentage change in DMR levels from pairwise comparisons: Pseudo *vs* Naïve, Trained *vs* Naïve, and Trained *vs* Pseudo. Percentage changes in methylation values are visualized on a color scale: blue indicates hypermethylation, and red indicates hypomethylation.

To identify methylation signatures specific to OD rule learning, we annotated differentially methylated regions (DMRs) and compared trained and pseudo-trained animals (**Fig. 4d**). The pseudo-trained group exhibited a distinct DMR profile (933 genes) enriched for pathways related to protein degradation, insulin signaling, and DNA methylation itself—including hypermethylation of *Dnmt3l*, *Smchd1*, and *Ehmt2*, suggesting broad epigenetic repression (**Suppl. Fig. 6a-c**). In contrast, trained animals showed unique hypomethylation at genes central to transcriptional regulation, chromatin remodeling, and synaptic plasticity, including *Grin1/2c/2d*, *Nr3c1*, *Jund*, *Fos*, and *Reln* (1145 genes) (**Fig. 4d and Suppl. Fig. 6d-f**). Interestingly, a shared set of 2515 DMRs between the two groups clustered into three major categories: (i) epigenetic gene regulation, (ii) mRNA transport and localization, and (iii) modulation of PKC pathways and the M-current (**Fig. 4d-g and** see pathway of (iii) in **Suppl. Fig. 2**). Yet, methylation patterns within these shared regions diverged markedly. For example, *Tet2*, a key demethylation enzyme, was hypermethylated in pseudo-trained but hypomethylated in trained animals—suggesting learning may activate a genome-wide demethylation program (e.g., ^29^). Similarly, genes linked to dendritic mRNA trafficking (*Eef1a1*, *Limk1*, *Arc*) were consistently hypomethylated in trained animals but hypermethylated in pseudo-trained counterparts (**Fig. 4g**). Strikingly, we observed hypomethylation in trained animals at key regulatory genes directly involved in the PKC signaling cascade and M-current modulation—including *Kcnq3*, *Kcnq4*, *Kcnq5* (which encode Kv7 channels), *Chrm4/5* (muscarinic receptors), *Gnaq*, *Gnas*, and multiple isoforms of phospholipase C (*Plcb2*, *Plcb4*, *Plcd3*, *Plcl2*), as well as *Akap5* (a PKC scaffolding protein). Hypomethylation at these loci suggests increased transcriptional accessibility, particularly of PKC, which can dominantly suppress the M-current regulatory pathway— potentially leading to enhanced intrinsic excitability and biophysical plasticity. Notably, the persistent reduction in M-current observed electrophysiologically in both F0 trained animals and their F1 offspring closely mirrors the epigenetic reprogramming identified in the paternal methylomes.

### Consistent pattern of altered DNA methylation in the hippocampus of trained F0 rats and their F1 offspring

We next examined whether methylation patterns in the hippocampi of F0 fathers were reflected in their untrained F1 offspring. Similar to their trained parents, F1 offspring of trained rats showed a pronounced bias toward hypomethylation compared to both naïve and pseudo-trained groups (**Figs. 5a-c**). In contrast, F1 offspring of pseudo-trained animals displayed a more mixed methylation profile, with substantial divergence from their parents in both the number and direction of methylation changes (see above). We next examined the intergenerational overlap of hypo- and hypermethylated regions (DMRs) and their associated genes across the hippocampal genomes of F0 rats and their F1 offspring (**Fig. 5d-f**) and quantified these relationships using GeneOverlap analysis (**Fig. 5g and see methods**). The most striking finding was that trained animals showed a substantially higher intergenerational overlap of hypo-DMRs with their offspring compared to other groups (**Fig. 5d-f bottom panels**). In contrast, pseudo-trained animals shared the largest and primarily hypermethylated overlap with their F1s (**Fig. 5d-f top panels**). Remarkably, the overlapping hypo-DMRs between trained F0 and F1 animals included key regulators of the M-current pathway, such as *Akap5*, *Kcnq5*, *Chrm5*, *Gsk3b*, and *Gnas* (**Suppl. Table 1**).

**Figure 5.**
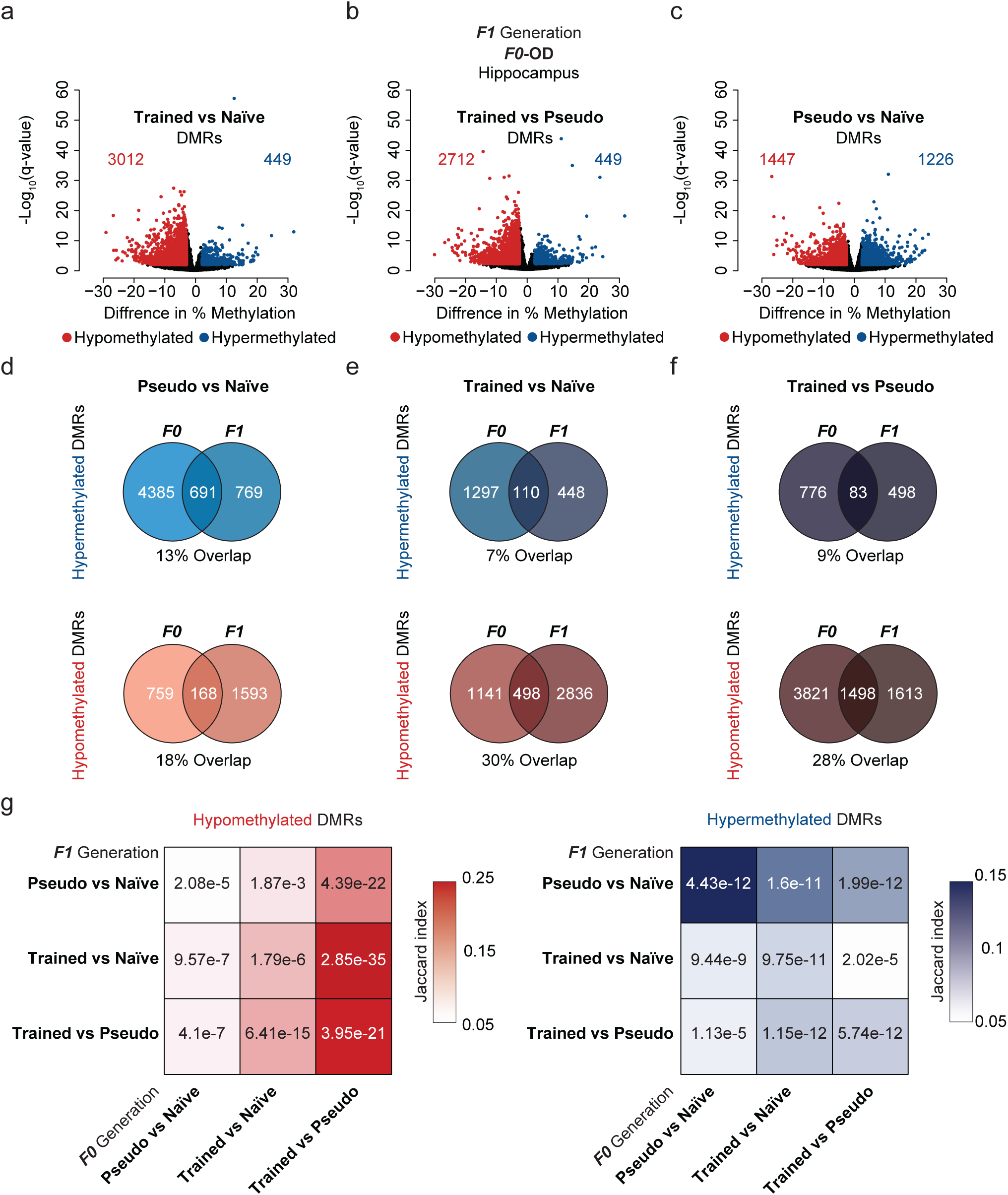
Superb learning capabilities are associated with consistent patterns of altered methylome in the hippocampus of both trained F0 rats and their subsequent F1 offspring. **(a-c)** Volacno plots of F1 hippocampal DMRs. Values of methylation from pairwise comparisons (difference in % methylation) are plotted against the average log_10_(p-value). Differentially hypo-methylated DMRs are shown in red, hyper-methylated DMRs are in blue, and non-significantly methylated DMRs are shown in black (and in color-coded numbers). Cutoff criteria – percentage change in methylations > +/-2.5, FDR<0.01, *n* = 2/per group. **(d-f)** Venn diagrams showing overlap of annotated Hypomethylated and Hypermethylated DMRs across the hippocampal genomes of F0 rats and their F1 offspring (via InteractiVenn; see **methods**). **(g)** Correlation analysis of annotated DMRs between F0 rats and their F1 offspring. P-value are presented in numbers and Jaccard index in color scale from Fisher’s exact were calculated by GeneOverlap R package and presented in the heatmap.

### Consistent patterns of altered DNA methylation in the hippocampus and sperm of F0 mice

Given the robust and consistent overlap of hypomethylation in key M-current– related genes across both trained fathers and their offspring, we next examined whether these epigenetic signatures are also associated with epigenetic changes in sperm^52^. By analyzing sperm methylation, we aimed to identify a stable, endogenous mechanism of transgenerational inheritance that preserves the complexity and specificity of the learning-induced epigenetic program.

To directly investigate germline transmission of epigenetic signatures linked to learning, we collected and purified sperm from F0 males during IVF and performed RRBS methylome profiling (cutoff: >2.5% methylation change, FDR <0.01). While hippocampal methylation profiles of trained and pseudo groups displayed clearly divergent hypo- and hypermethylation patterns, their sperm methylomes showed markedly fewer DMRs (compared to naïve) and lacked pronounced global directional differences (**Figs. 6a-c**). However, annotation analysis revealed that, unlike in the hippocampus, DMRs in sperm from both trained and pseudo groups were significantly enriched in non-coding RNA (ncRNA) species—including lncRNAs, miRNAs, snoRNAs, and snRNAs (**Fig. 6d**). When separating sperm DMRs into coding and non-coding loci, we found that both groups also harbored methylation changes in protein-coding genes associated with neuronal processes (**Suppl. Fig. 7**). Notably, gene enrichment in trained animals was particularly biased toward pathways governing learning, memory, hippocampal signaling, and exploratory behavior. Furthermore, a strong presence of DMRs in genes regulating the cAMP signaling pathway aligned with hippocampal methylation data (see **Suppl. Fig. 6**), reinforcing the hypothesis that sperm-borne epigenetic modifications, especially in regulatory ncRNA and learning-relevant genes, may serve as a mechanistic substrate for inherited learning capacity.

**Figure 6.**
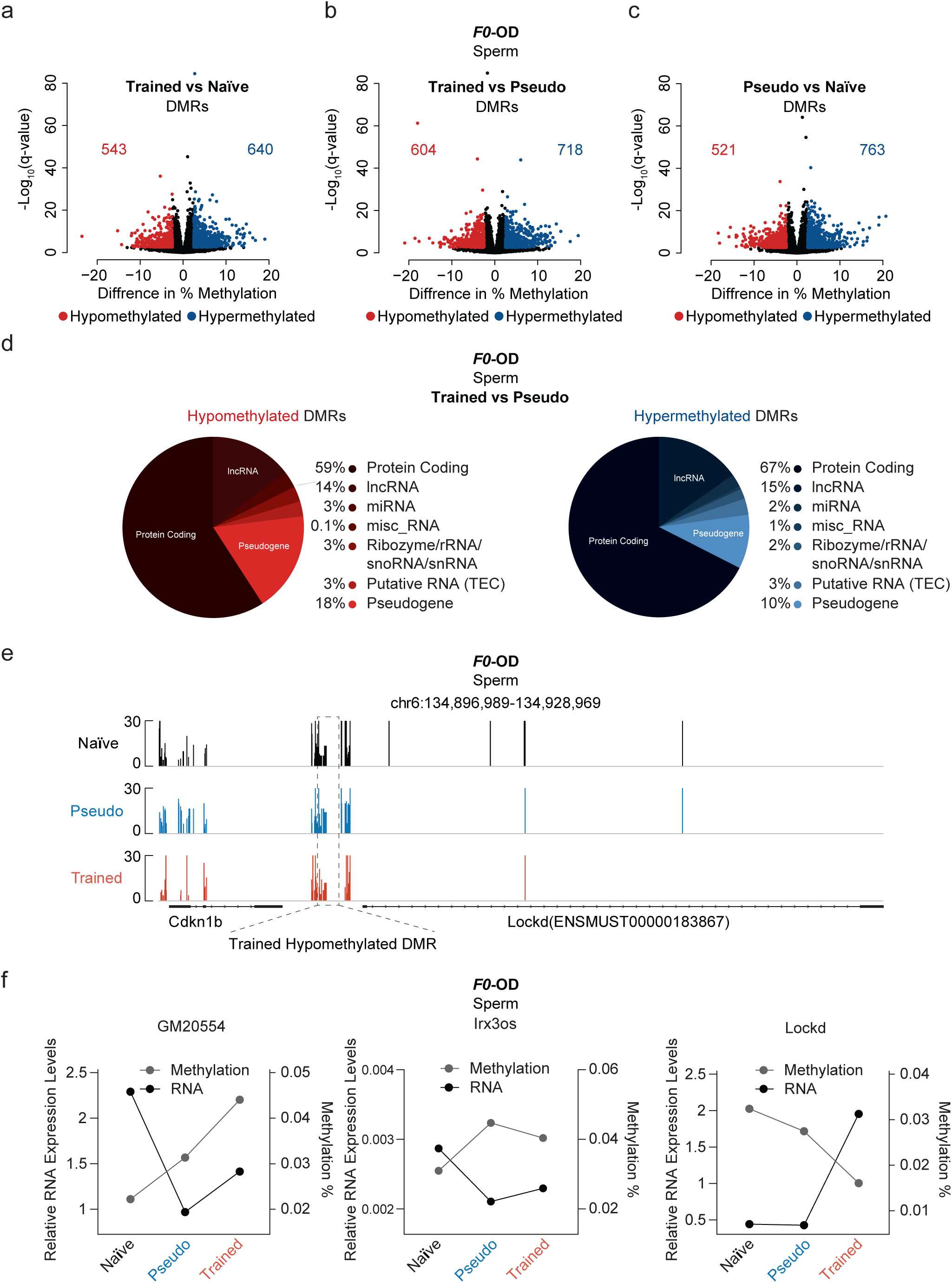
Training of F0 males alters their sperm methylation patterns, predominantly around genomic regions encoding for ncRNA and modulation of the M-current. **(a-c)** Volcano plots for sperm DMRs. Values of methylation from pairwise comparisons (difference in % methylation) are plotted against the average log_10_(P value). Differentially hypomethylated DMRs are shown in red, differentially hypermethylated DMRs are in blue, and non-significantly methylated DMRs are shown in black (and in color-coded numbers). Cutoff criteria – percentage change in methylations > +/-2.5, FDR<0.01, *n* = 5-6/per group. **(d)** Pie chart showing genomic distribution of Hypomethylated and Hypermethylated DMRs to protein coding regions and various non-coding RNA species. **(e)** Representative methylation plots showing hypomethylated DMR identified in the sperm of F0 trained mice near the ENSMUST00000183867 ncRNA locus. The IGV genome browser plot presents the average percentage of methylation levels, with DMRs highlighted by red rectangles. (**f**) Plot showing percent methylation in gray and relative mRNA expression levels in black of the ncRNAs *Gm20554*, *Lockd*, and *5033426O07Rik*. Average percent methylation per group was calculated from the RRBS dataset. RT-qPCR was performed using designated primers on pooled sperm samples (n=5-6 per group). mRNA expression levels are presented relative to β-Actin.

Building on our behavioral, physiological, and epigenetic evidence implicating the M-current pathway in transgenerational learning, we next focused specifically on sperm-DMRs located within genomic regions encoding for ncRNAs, which may regulate this pathway. Using a curated list of M-current–related genes (**Fig. 4g**), we performed *in silico* analysis to identify ncRNAs predicted to regulate their expression. We then intersected this set with the ncRNAs found to harbor DMRs in sperm samples (**Table 1**). Strikingly, while only two DMRs on ncRNAs were common between trained and pseudo groups, these were largely linked to general regulators such as PLC (*Gsk3b* and *Plcb2*), which influence multiple signaling cascades and lack specificity for M-current modulation. In contrast, trained animals exhibited several unique DMRs on ncRNAs, including hypermethylation of *ENSMUST00000177421 (Gm20554), ENSMUST00000212635 (5033426O07Rik)* and hypomethylation *ENSMUST00000210265 (Plcb4), ENSMUST00000183867 (Lockd), ENSMUST00000222673 (Gm34923)* (**Table 1 and Fig. 6e**). These ncRNAs were predicted to regulate key components of the M-current pathway—including *Chrm5*, *Akap5*, *Calm1*, and *Kcnq5* (**Table 1**). Consistent with the observed hypermethylation patterns (gray line), trained animals showed reduced relative RNA expression levels (black line) of *ENSMUST00000177421* (*Gm20554*) and *ENSMUST00000212635* (*5033426O07Rik*) in sperm compared to naïve controls (Fig. 6f). Additionally, expression of *ENSMUST00000183867* (*Lockd*) was also decreased, corresponding to the hypermethylation detected at its genomic locus in trained animals relative to controls (**Fig. 6f**). These strongly suggest that training induces targeted epigenetic changes in sperm ncRNAs that are specifically poised to regulate key nodes in the M-current signaling pathway. Together with our earlier data, these findings support a mechanistic model in which sperm-borne ncRNAs mediate the inheritance of enhanced learning by modulating the same molecular axis—M-current suppression—identified functionally in the offspring.

**Table 1.** Putative Regulative ncRNA Found Significantly Methylated in Sperm.

Collectively, our data show that rule learning induces substantial alterations in the hippocampus of trained rats and sperm samples of trained mice. The parallel changes in M-current regulation observed across two species (mouse and rat), spanning brain and sperm methylation patterns, as well as biophysical and behavioral modifications across two generations, reinforce the notion that this represents a conserved mechanism of epigenetic inheritance.

## Discussion

### Transgenerational Epigenetic Inheritance of Rule Learning: A Novel Neurobiological Framework

A core principle of epigenetics is that salient experiences—especially those engaging emotional, cognitive, and behavioral processes—can leave molecular imprints in the brain. These experience-dependent marks, can durably reprogram gene expression and neural plasticity, ultimately shaping behavior. Growing evidence now extends this framework to the germline, showing that the same experiences can induce complementary epigenetic modifications in sperm and oocytes^1–9^. These changes may arise through brain-to-gonad signaling that remains to be defined, or through parallel alterations established directly in germ cells. Such germline marks—typically shifts in DNA methylation or ncRNA profiles—can be stably inherited, reprogramming brain function or development and shaping behavior in successive generations. Yet, to date, most mammalian studies on transgenerational epigenetic inheritance have focused on simple behavior or learning paradigms— typically those involving aversive conditioning, such as fear responses or avoidance behaviors^1,3,8,17^. Whether more complex cognitive processes, such as *rule learning*— the ability to extract and generalize abstract principles across contexts—can be inherited epigenetically has remained unexplored.

In the present work, we provide compelling evidence for the transgenerational epigenetic inheritance of a high-order cognitive function known as rule learning, or the ability to “learn how to learn”. Our collection of findings show:

1. Inheritance of Complex, Generalized Learning Abilities. Offspring of F0-trained animals—up to the F3 generation—exhibited exceptional learning capabilities even without exposure to training or reinforcement (**Fig. 1b-e**). Unlike previous models limited to specific cues or threats, the enhanced cognitive performance in our paradigm generalized across distinct tasks, sensory modalities, and behavioral demands (**Fig. 1a and Suppl. 5**), highlighting a *global learning state* rather than task-specific memory.
2. Robust Epigenetic Transmission Beyond Parental Behavior. Cross-fostering paradigms, in which offspring of trained and untrained mothers were reared by surrogate mothers from the opposite group, ruled out social learning as a mechanism (**Fig. 1f**). Moreover, offspring from naïve females fertilized with sperm from trained males also exhibited superior learning, establishing that the inheritance is epigenetic and not maternally conveyed through care or environment (**Fig. 1h**).
3. Neuronal Excitability as a Cellular Signature of Inherited Learning. Functional analyses revealed that neurons from the offspring of trained animals displayed enhanced intrinsic excitability (**Fig. 2**)—akin to changes that have been shown to occur in neurons of trained animals (in the ancestral F0 generation) following intense cognitive training^20–22^. Specifically, hippocampal (CA1, CA3), mPFC, and BLA neurons showed significantly reduced medium and slow afterhyperpolarizations (AHPs), facilitating increased spike output during learning (**Fig. 2a-d and Suppl. Fig 4**).
4. M-current Downregulation via Persistent PKC Activation as driver of excitability. Inherited changes were mechanistically linked to a persistent downregulation of the cholinergic M-current (**Fig. 3**), a potassium current known to regulate neuronal excitability and plasticity. In F1 offspring, we found that the medium and slow AHP reductions were reversed by blocking PKC, implicating long-term PKC-dependent modulation by at least two converging pathways (**Fig. 3 and Suppl. Fig. 3**). This precisely mirrors the M-current suppression observed in rule-learned F0 parents^20–22,26,44–46^.
5. Causal Relationship Between M-current and Cognitive Performance. Pharmacological manipulations *in vivo* further confirmed the causal role of M-current modulation. Scopolamine, a muscarinic receptor antagonist, impaired rule learning in controls but had no effect in offspring of trained rats, suggesting the M-current was already suppressed in the latter. Conversely, XE991, a KCNQ channel blocker, enhanced learning in controls but had no effect in offspring of trained animals—equalizing learning performance between the groups (**Fig. 3j-l**).

### Epigenetic Underpinnings: Methylation and Non-Coding RNAs

Our molecular data further support the transgenerational nature of these effects. Whole-genome methylation analyses revealed substantial and directionally distinct methylation changes in the hippocampi of trained F0 rats and their F1 progeny (**Fig. 4 and 5**). While pseudo-exposed animals exhibited largely hypermethylated regions, trained animals displayed focused hypomethylation (**Fig. 4a-c**), particularly in genes involved in M-current regulation and PKC signaling (**Fig. 4e-g**). Strikingly, these same hypo-DMRs were found in naïve F1 offspring of trained fathers—indicating that the methylome itself may carry cognitive capacity instructions (**Fig. 5**).

Interestingly, sperm from trained F0 males lacked the gene-associated methylation pattern we observed in hippocampus (**Fig. 6a-c**). Instead, differential methylation clustered on non-coding RNAs (ncRNAs), particularly long non-coding RNAs (lncRNAs) predicted to regulate M-current-related genes (e.g., *Charm5, Akap5, Calm1, Kcnq5*) (**Fig. 6d-e**). Notably, these methylation changes coincided with parallel alterations in the expression levels of the same lncRNAs, reinforcing their potential role in transmitting M-current–related regulatory information to the next generation (**Fig. 6f**). Although the mechanism establishing these gonadal marks remains unclear, the data point to a bifurcated model: the brain stores functional methylation marks, whereas sperm conveys a regulatory blueprint via dual methylation and ncRNA-mediated mechanisms —an idea consistent with emerging models of sperm-mediated intergenerational transmission^53–55^.

Moreover, the observation that even F₂ and F₃ descendants of trained males display heightened neuronal excitability and superior learning supports the notion that the brain can relay information to sperm and that germline epigenetic changes are reinforced across multiple learning bouts rather than arising from a single event. Establishing the precise mechanisms of this soma-to-germline communication remains an important goal for future studies.

### A Dual-Layer Model for Inherited Rule Learning

We propose a novel, dual-layer epigenetic model in which rule learning initiates parallel molecular signatures in both the somatic (brain) and germline (sperm) tissues:

1. In the Brain: Enhanced learning induces stable hypomethylation and M-current suppression via PKC activation, directly augmenting neuronal excitability and plasticity.
2. In the Germline: Epigenetic reprogramming is mediated primarily by differentially methylated ncRNAs that regulate the same cognitive pathways, providing a mobile blueprint for transmission.

This integrated system enables the stable and specific transmission of a complex cognitive phenotype across generations and challenges the notion that only maladaptive or simple behaviors are epigenetically inherited.

Together, our findings fundamentally shift the landscape of epigenetic inheritance. We demonstrate that mammals can transmit abstract cognitive strategies through precise molecular mechanisms. This challenges long-held boundaries between nature and nurture, opening new avenues for research into learning, memory, psychiatric resilience, and even educational interventions informed by inherited neuroplasticity.

## Methods

### Complex olfactory-discrimination learning

Subjects were male and female rats and mice aged 2 months. Prior to training they were maintained on a 23.5 hr. water deprivation schedule with food available ad libitum.

#### Apparatus and odors

Olfactory discrimination protocol was performed in a 4-arm radial maze, with odors commonly used in the food industry (**Fig. 1a**).

#### Training

Olfactory training will consist of 20 trials per day as previously described^20,27^. Learning was considered as acquired upon demonstration of at least 80% positive-cue choices for the last 10 trials of the day. A pseudo-trained group of age-matched rats was exposed to the same protocol of training, but with random water rewarding. An age-matched naive group of rats were water-deprived but was not exposed to any training. When trained in this complex task, rats demonstrate increased capability to discriminate between new odors once they reach good performance with the first pair of odors^20,56^. Thus, two learning phases are clearly apparent: The first phase of learning the rule, which usually requires 7-8 days, in which rats develop a strategy for performing the task, and the second phase of enhanced learning capability after rule-learning, when learning new odors requires 1-2 training days. Pseudo-trained rats did not show a preference for any odor throughout the training period. Mice which are trained in similar, but smaller maze, show the same learning capabilities.

### Water Maze Training

Water maze training was performed as previously described^26^. Animals were trained in a circular pool 1.8 m in diameter and 0.6 m high, containing water at 26 ± 1°C. The pool will occupy the center of a room and will contain various salient cues. A 10 cm^2^ transparent platform was hidden in a constant position in the pool with its surface submerged 1 cm below the water level. Animals were given four consecutive training sessions per day, starting from random locations around the pool each time. Each session will last up to 60s. If the animal did not reach the platform within this time, it was guided to it. The animal was allowed to remain on the platform for 30 s to learn its location. Each animal was trained for 4 days. The latency time to reach the platform was measured by using fixed overhead video recordings, analyzed offline.

### Auditory Fear Conditioning

Auditory fear conditioning was performed as previously described^57^. Rats were trained in a Plexiglas rodent conditioning chamber with a metal grid floor. The animals were habituated to the context on the first two days for 10 minutes, followed by conditioning training day; where the animal was given 5 mins of acclimation to the chapter followed by five pairings of a tone (parameters-CS:30s,4 kHz, 80 dB) accompanied by a foot shock (parameters-US:1 s, 0.5 mA). The inter-trial interval was two minutes. Extinction was examined for three days following the conditioning day, in which animals were given five minutes for accumulation to the chamber followed by five tones without the foot shock. The total time of all the days was 15 mins and freezing levels were measured the whole period.

### Slice preparation and recordings

Coronal brain slices, 400 µm prepared from the hippocampus from rats aged 1-2 months, BLA and mPFC as previously described^58^ fand kept in oxygenated (95% O2 + 5% CO2) Ringer’s solution containing (in mM): NaCl, 124; KCl, 3; MgSO_4_, 2; NaH_2_PO_4_, 1.25; NaHCO_3_, 26; CaCl_2_, 2; and glucose, 10. Intracellular recordings were performed at 34.5 °C, with 4 M K-acetate filled sharp glass microelectrodes with an axoclamp 2A amplifier and analyzed using pCLAMP software. The identity of animals (paradigm and day of training) from which the neurons were taken and recorded was not known to the person conducting the experiments and measurements.

#### Synaptic Stimulation

Stimulation was done by stimulating the Schaffer Collateral 20 times at 50 Hz. Stimulation amplitude was adjusted such as the EPSP of the recorded pyramidal neuron was 10 mV as previously described^59^.

#### Membrane properties

Membrane properties were measured in current clamp recordings. Input resistance was determined by linear regression fit to a voltage/current curve. Membrane time constant will be determined by fitting one exponential curve to the membrane voltage decay at the end of a 10 mV, 200 ms long hyperpolarizing step. Spike amplitude was measured from the resting potential to the peak. Spike width was measured at the threshold.

#### post-burst AHP measurements

For post-burst AHP measurement, neurons were depolarized in current clamp mode to Vm of -60 mV with DC current injection, and 100 ms-long depolarizing current steps was applied with intensity sufficient to evoke trains of 6 action potentials. The medium and late AHPs amplitudes were measured from baseline to the peak of the hyperpolarizing voltage deflection that follows the spike train (see figure 2a for example). The AHP value was determined from the averaged amplitude in 4-5 consecutive traces, evoked at intervals of 10 sec.

### In Vitro Fertilization (IVF) procedure

IVF was performed using mice sperm from mice of the three groups (17 mice total, 6 naïve, 6 pseudo-trained, 5 trained) through the complex olfactory-discrimination task. F0 males were trained or not in the complex olfactory-discrimination task and 10d after, sperm was collected from the cauda epididymis. The sperm was cryopreserved using a cryoprotective agent (CPA) supplemented with L-Glutamine. The sperm was sealed in cryo-straws, with 10 µL of sperm in each straw, and stored in liquid nitrogen. Sperm collection was conducted at the Cryopreservation and IVF Core of the Pre-Clinical Research Authority at the Technion - Israel Institute of Technology, Haifa, Israel. The cryopreservation protocol followed the European Mouse Mutant Archive (EMMA) guidelines (https://www.infrafrontier.eu/emma/cryopreservation-protocols, Sperm freezing using CPA that has been supplemented with L-Glutamine). Female wild-type (WT) C57Bl/6JOlaHsd mice, 4 weeks old, unrelated to the experimental groups, were used as oocyte (egg) donors. The female mice were hormonally stimulated to induce superovulation (multiple oocytes in one cycle). This was achieved by administering pregnant mare serum gonadotropin (PMSG) to induce follicular growth, followed by human chorionic gonadotropin (hCG) to trigger ovulation. The superovulated 4-week-old females were sacrificed 12-14 hours after the hCG injection to collect the oocytes for IVF. Oocytes were placed in a GSH HTF culture medium under a microscope. A sperm straw was thawed and placed in TYH+MBCD medium to enhance sperm function. The equilibrated sperm was added to the oocyte plate for fertilization. Four hours post-fertilization, the embryos were transferred to one of four HTF drops for development into the next stages of embryos. Twelve hours post-fertilization, the cultured 2-cell stage embryos were implanted into the infundibulum of pseudo pregnant foster ICR female mice, 8 weeks old. 9-11 embryos were transferred into two oviducts of each female. To induce pseudopregnancy, the ICR female was mated with a vasectomized male for verification of estrus stage. Pregnancy is confirmed by checking for the presence of embryos in the foster mother’s uterus, usually 7-10 days after the embryo transfer. If successful, the foster mother will carry the embryos to term and give birth to the pups. Pups were weaned from their foster moms at 3.5 weeks of age and reared to 1-2 months of age before training in the complex OD task. IVF process was conducted at the Cryopreservation and IVF Core of the Pre-Clinical Research Authority at the Technion - Israel Institute of Technology, Haifa, Israel. The IVF protocol followed the European Mouse Mutant Archive (EMMA) guidelines (IVF recovery procedure using freshly harvested sperm, incorporating methyl-β-cyclodextrin and reduced glutathione).

### Animals Ethics

This study was carried out in strict accordance with the Guide for the Care and Use of Laboratory Animals of the National Institutes of Health. All experimental procedures and protocols were approved by the ethical committee of the University of Haifa for experimentation with animals and were performed in strict accordance with University of Haifa animal ethical regulations. (Permit number, UoH - IL - 2203 - 138 - 3).

### DNA Methylation

#### Hippocampus

12 rats – 4 rats for each group naïve, pseudo-trained and trained. For each group, there were 2 father-son pairs (F0 and F1) meaning that that for each of the 6 groups (F0 naïve, F0 pseudo-trained, F0 Trained, F1 naïve, F1 pseudo-trained, F1 Trained) there were 2 animals. It is important to note that F1 animals were not exposed to any training or task. F0 were about 4 months old and F1 were about 1 month old. Hippocampi were extracted and prepared loosely as described^60^. Briefly – rats were euthanized, hippocampi were extracted and dissociated by trypsin and manual triturations and strained with 40 µm strainers. Cells were pooled by group, then collected and pelleted, then frozen overnight in -80°C in PBS. Genomic DNA was extracted by DNeasy Blood and Tissue (Qiagen, cat no. 69504). DNA purity was assessed using Nanodrop, according to the 160/280 absorbance ratio. All samples showed a ratio of 1.8-2.0, indicating good purity. DNA samples were quantified using Qubit 2 (Thermo Fisher) using the Qubit dsDNA HS Assay Kit (Invitrogen, cat no. Q32854). 6 RRBS libraries were constructed simultaneously using Zymo-Seq RRBS Library Kit (Zymo Research, cat no. D5460) according to the manufacturer’s protocol. 50 ng DNA was used as starting material. After construction, the concentration of each library was measured using the Qubit dsDNA HS Assay Kit (Invitrogen, cat no. Q32854) and the size was determined using the TapeStation 4200 (Agilent) with the High Sensitivity D1000 kit (cat no. 5067-5584). All libraries were mixed into a single tube with equal molarity. The sequencing data was generated on Illumina NextSeq2000, using P3 100 cycles (Read1-51; Index1-8; Index2-8; Read2-51) (Illumina, cat no. 20040561). For the bioinformatics analysis the quality control was assessed using FastQC (v0.11.8). Reads were trimmed for adapters, minimum quality Phred score of 20 and a minimum length of 20 using trim_galore (v0.6.5) with specific flags for RRBS libraries: “--rrbs --non_direction.a”l Trimmed paired reads were aligned to Rattus norvegicus (Rnor_6.0) reference genome using two bisulfite sequencing (WGBS/RRBS) directed aligners: Bismark (v0.24.0) with default parameter settings (“--non_directional” flag was used). Following alignment with Bismark, methylation extraction was carried out with “bismark_methylation_extractor” script (built-in to Bismark software). By manual review of methylation bias plots, additional trimming of 4 bases on both 5’ and 3’ ends was performed for both read #1 and #2 to remove noise. Following methylation extraction, Bismark outputs were read into R and analyzed for DMRs and differential methylation using methylKit package (v1.20.0). Features with less than 3 coverage each were filtered out. Additionally, the top 0.05% of covered positions were removed. To address the multiple comparisons issue, FDR was estimated using the q-value method. DMRs were calculated using window and step size of 1000 with minimum number of bases per DMR of 10. Quantified positions and regions were annotated using “genomation” package (v1.26.0). Further analysis was done using custom R code, Metascape (https://metascape.org/), InteractiVenn (https://www.interactivenn.net/), Revigo (http://revigo.irb.hr/), and additional visualizations were done using Integrative Genomics Viewer (https://igv.org/). DNA QC, RRBS library preparation and NGS were conducted by the Technion Genomics Center, Technion-Israel Institute of Technology, Haifa, Israel. The bioinformatic analysis was conducted in collaboration with the Technion Genomics Center.

#### Sperm

Sperm obtained from the IVF procedure from 6 Naïve F0, 6 Pseudo-trained F0 and 5 Trained F0 mice were pooled into groups of 3 (2 pools per group, in trained a group of 3 and a group of 2). Genomic extraction for the sperm was done twice with varying amount of sperm. Each pool of sperm was done separately, meaning in total 12 separate methylation runs. Sperm genomic DNA extraction was adapted from previous protocols^61^. In brief, since our sperm sample was already purified when sperm extraction was done for the IVF procedure, we started at step 2 of the protocol^30^ adding the sperm pellet to 450 µl of buffer RLT+ (Qiagen, cat no. 1053393) and 50 µl of TCEP (Thermo Scientific, cat no. 77720). We then added 0.1g of stainless-steel bashing beads (NextAdvance, cat no. ADV-SSB02-RNA) and bash for 5 mins with a strong vortex. Immediately after genomic DNA was extracted using the provided standard protocol in the AllPrep DNA/RNA Mini Kit (Qiagen, cat no. 80204). DNA samples were quantified using Qubit 4 (Thermo Fisher) using the Equalbit dsDNA HS Assay (Vazyme, cat no. EQ121). 12 RRBS libraries were constructed simultaneously using Zymo-Seq RRBS Library Kit (Zymo Research, cat no. D5460) according to the manufacturer’s protocol. 50ng DNA was used as starting material. After construction, the concentration of each library was measured using the Equalbit dsDNA HS Assay (Vazyme, cat no. EQ121) and the size was determined using the TapeStation 4200 (Agilent) with the High Sensitivity D1000 kit (cat no. 5067-5584). All libraries were mixed into a single tube with equal molarity. The sequencing data was generated on Illumina NextSeq2000, using P2 100 cycles (Read1-51; Index1-8; Index2-8; Read2-51) (Illumina, cat no. 20046811). For the bioinformatics analysis the quality control was assessed using FastQC (v0.12.1). Reads were trimmed for adapters, minimum quality Phred score of 20 and a minimum length of 20 using trim_galore (v0.6.10) with specific flags for RRBS libraries: “--rrbs --non_directional”. Trimmed paired reads were aligned to Mus musculus (GRCm39) reference genome using the bisulfite sequencing (WGBS/RRBS) directed aligner Bismark (v0.24.0) with “--non_directional” flag used and parameter settings set to default. Following alignment with Bismark, methylation extraction was carried out with “bismark_methylation_extractor” script (built-in to Bismark software). By manual review of methylation bias plots, additional trimming of 3 and 4 bases on 5’ and 3’ ends, respectively, was performed for both read #1 and read #2 to remove noise. Following methylation extraction, outputs were read into R and analyzed for DMRs using methylKit package (v1.20.0). Initially features with less than 10 coverage were filtered out. Additionally, the positions with coverage >5000 were removed (about 0.03% of covered positions). To find differentially methylated regions, regions were defined using a 1 kb sliding window over the genome. Genomic tiles containing at least 10 CpG positions, each with at least coverage of 3 reads, were defined as regions to be compared. To address the multiple comparisons issue, FDR was estimated using the q-value method. Quantified positions and regions were annotated using “genomation” package (v1.26.0) using genomic regions extracted from Mus musculus (GRCm39) reference genome and CpG isle and shore coordinates according to the UCSC Genome Browser CpG track (GRCm39) exported using the Table browser. Additionally, in order to test as many CpG and 1K tile regions as possible, while utilizing as much statistical power available, we merged samples with a minimum of 3 samples in each group and performed differential methylation tests on every shared CpG position and 1K tile. The tests performed include the tests of all samples and the additional positions where at least 3 samples of each treatment group have methylation data. This approach yielded a higher number of tests but did not increase dramatically the number of significantly differentially methylated positions found. Further analysis was done using custom R code, Metascape (https://metascape.org/), InteractiVenn (https://www.interactivenn.net/), Revigo (http://revigo.irb.hr/), and additional visualizations were done using Integrative Genomics Viewer (https://igv.org/). DNA QC, RRBS library preparation and NGS were conducted by the Technion Genomics Center, Technion-Israel Institute of Technology, Haifa, Israel. The bioinformatic analysis was conducted in collaboration with the Technion Genomics Center.

### RNA isolation, cDNA preparation and real-time polymerase chain reaction

Six sperm samples from each group was pulled together and RNA was isolated using Quick-RNA Miniprep Plus Kit (Zymo Research) according to the manufacturer’s instructions. RNA was reverse-transcribed to single-stranded cDNA by Super Script II Reverse Transcriptase (Thermo Fisher Scientific) and random primers. qPCR was performed in a CFX Connect Real-Time PCR Detection System (Bio-Rad) with PerfeCTa SYBR Green FastMix (Quanta BioSciences). Dissociation curves were analyzed following each real-time qPCR to confirm the presence of only one product and the absence of primer dimer formation. The threshold cycle number (Ct) for each tested gene (X) was used to quantify the relative abundance of that gene using the formula 2^-(Ct geneX – Ct standard)^. *β-Actin* was used as the standard for mRNA expression. Primers used for the real-time qPCR and gene specific amplification are listed below.

### Primer List

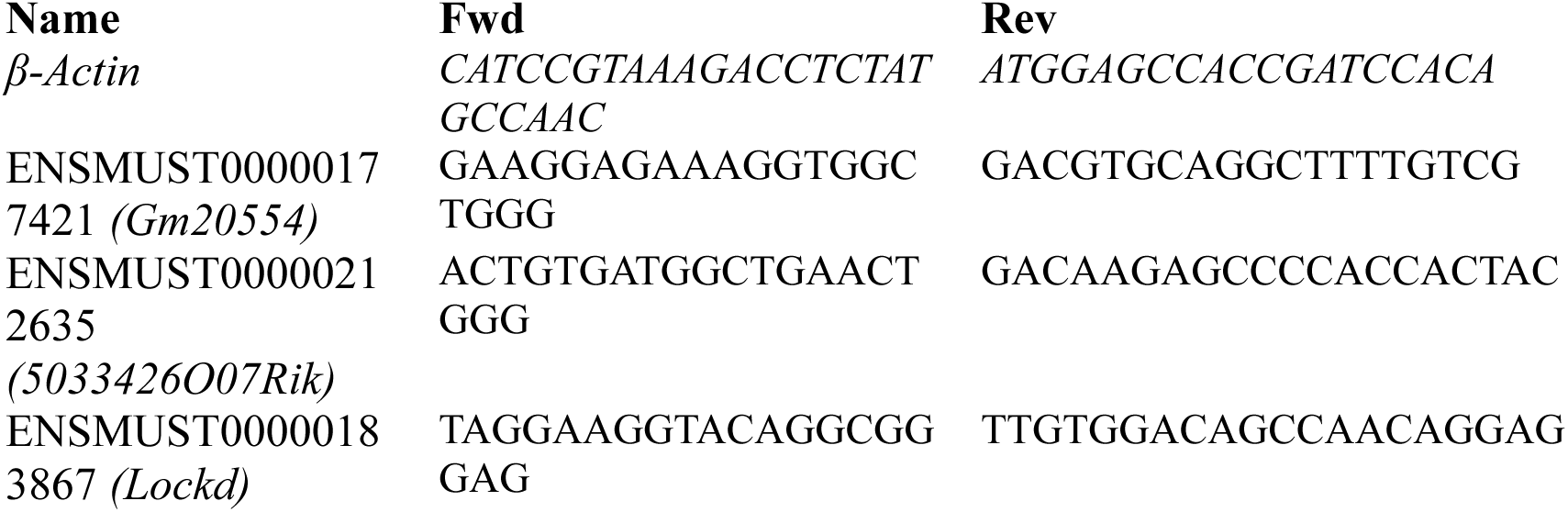

### In-Silico ncRNA Identification

To identify putative ncRNAs regulating the expression of genes associated with the M-current, we utilized the open-source LncRRIsearch tool, a web server for lncRNA-RNA interaction prediction integrated with tissue-specific expression and subcellular localization data^62^. Local base-pairing interactions between putative ncRNAs and target genes were analyzed. For downstream analysis, only lncRNAs enriched in both the brain and testis were selected.

### Statistical Analysis

All results are shown as mean ± SEM. multiple-group comparisons were performed using parametric and non-parametric (depending on normality) one-way analysis of variance (ANOVA) with either post hoc Dunns test or Dunnett’s multiple group comparison. Multiple group comparisons in different days were done by two-way ANOVA with Tukey’s test as post hoc. Two-group comparisons were performed using parametric and non-parametric (depending on normality) two-tailed t test, and pairwise comparisons were performed using a paired t test (GraphPad Prism 8). Linear regression was performed using simple linear regression. Data are presented as mean ± SEM. Significance is indicated as *, P<0.05; **, P<0.01; ***, p<0.001; n.s., non-significant.

## FIGURES LEGENDS

**Supplementary Figure 1.**
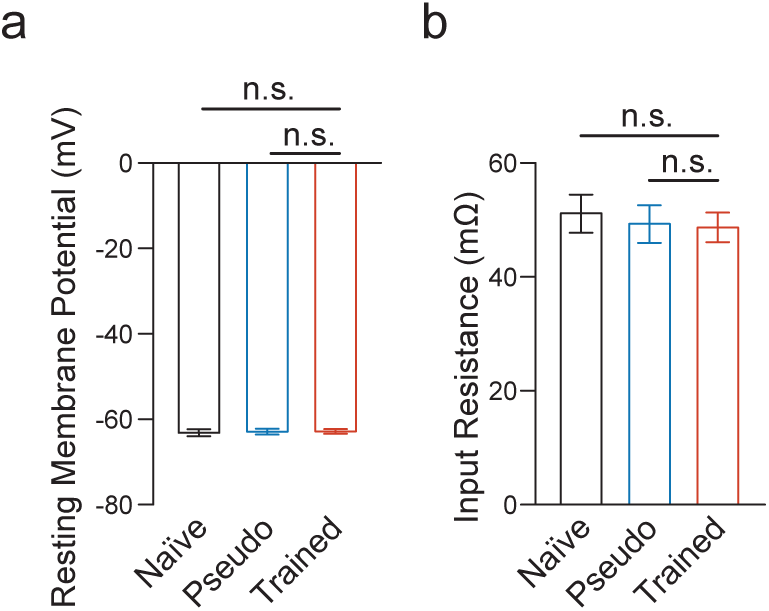
Membrane properties are not changed due to learning. **(a, b)** Resting membrane potential and input resistance are comparable across neurons from all three experimental groups Data are presented as mean ± SEM. *, P<0.05; **, P<0.01; ***, p<0.001; n.s., non-significant.

**Supplementary Figure 2.**
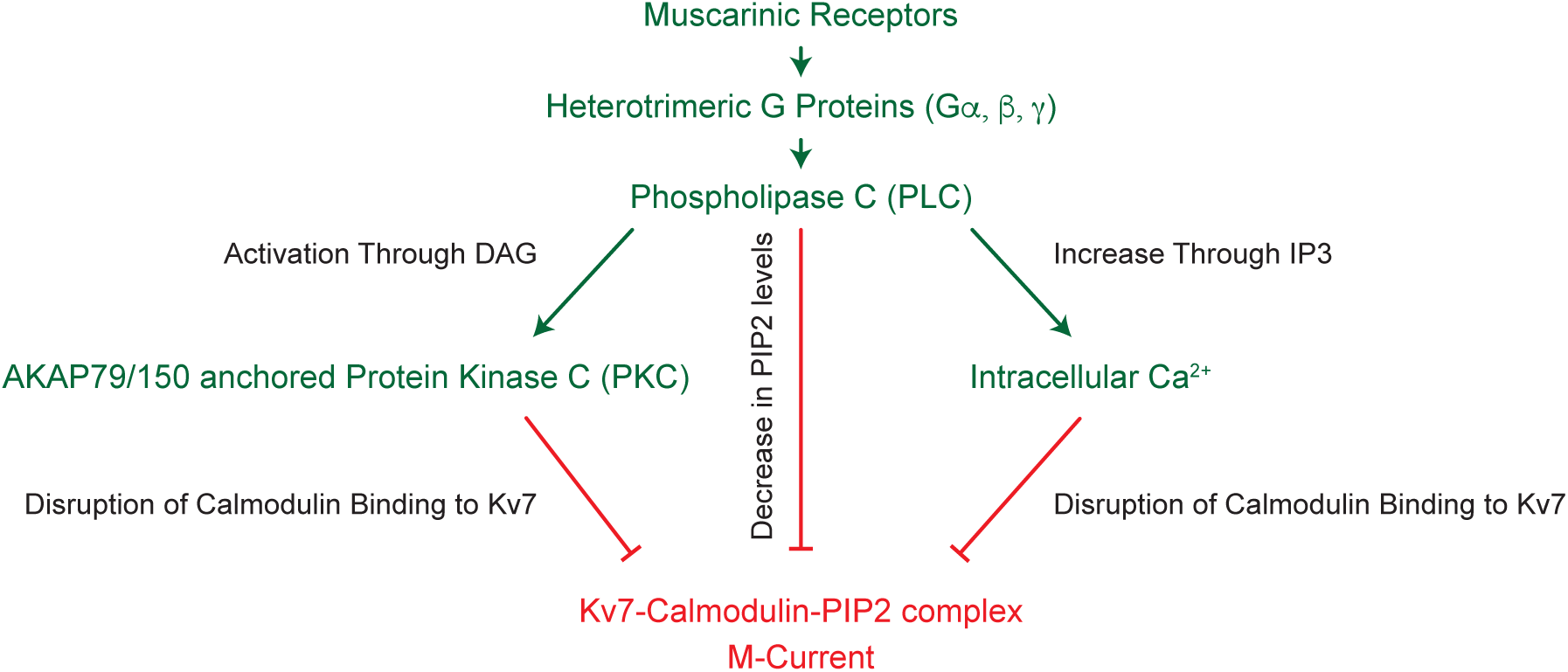
Signaling mechanism leading to suppression of the M-Current by. Activation of metabotropic muscarinic receptors activates G proteins leading to activation of PLCβ and subsequent decrease in PIP2 levels, production of inositol-1,4,5-trisphosphate (IP3) and diacylglycerol (DAG). IP3 activates IP3 receptors (IP3R) at the ER to elevate intracellular calcium (Ca^2+^), whereas DAG activates PKC. These three pathways converge to inhibit the KCNQ (Kv7)-current (M-current) (adapted from^40–43)^.

**Supplementary Figure 3.**
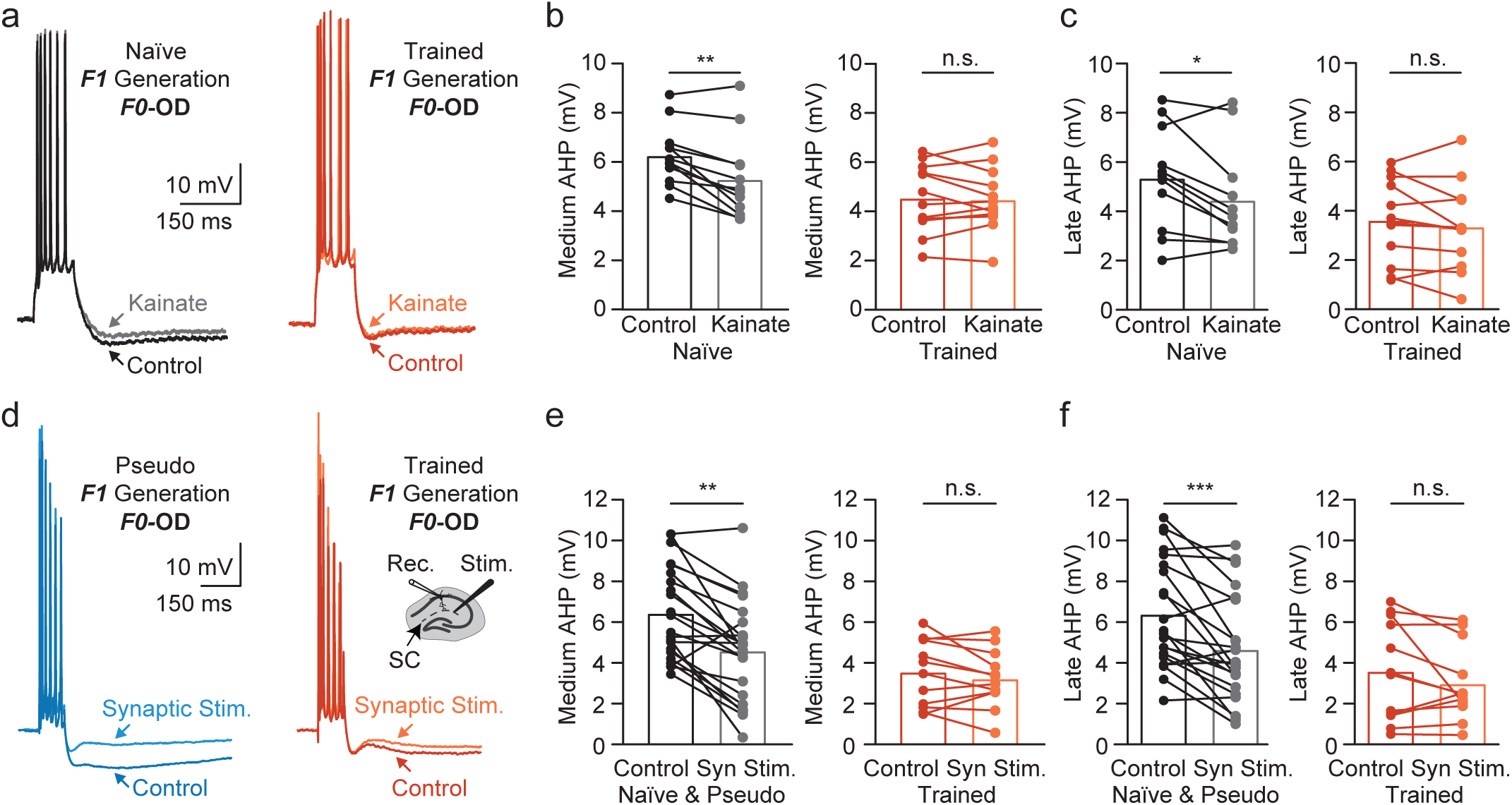
Reduced AHP in neurons F1-progenies from trained rats is maintained by persistent GluK2-activation. **(a)** Representative traces from neurons from progenies of F0-Naïve (black) or F0-Trained parents (orange). Application of kainate reduces the amplitude of the medium and late AHP in F1 from naïve animals (grey trace), with no effect in F1 neurons from Trained animals (light orange), summarized in **(b)** and **(c)**, respectively. **(d)** Two representative examples of the effect of synaptic stimulation on the post-burst AHPs evoked in a neuron from a pseudo-trained rat offspring (cyan/blue) and a trained rats’ offspring (orange/red). *Inset*: location of stimulating and recording electrodes. **(e)** Synaptic stimulation reduces the amplitude of the medium and late AHP in F1 progenies from naïve or pseudo-trained animals, with no effect in F1 neurons from Trained animals, summarized in **(e)** and **(f)**, respectively. Data are presented as mean ± SEM. *, P<0.05; **, P<0.01; ***, p<0.001; n.s., non-significant.

**Supplementary Figure 4.**
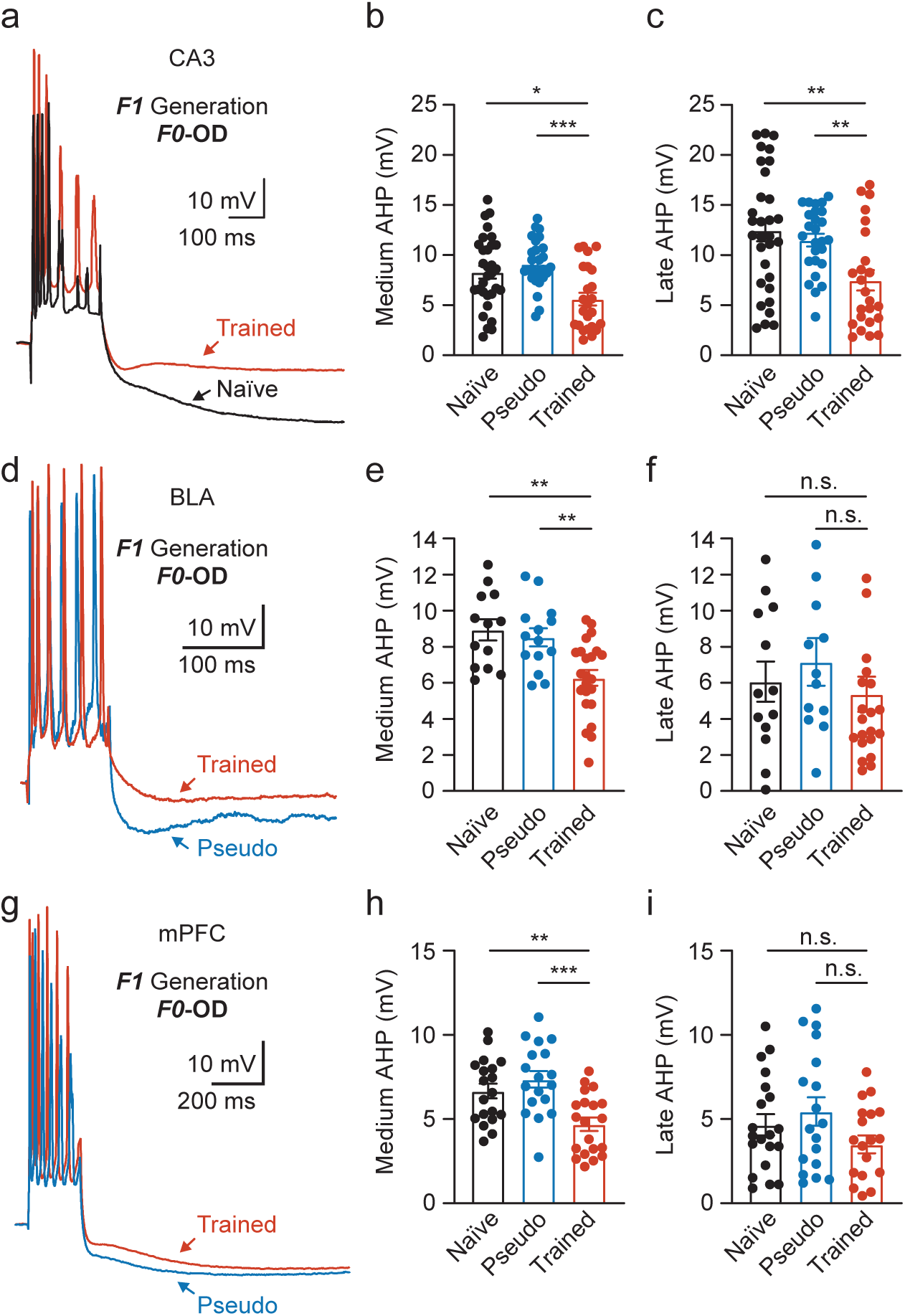
Inherited enhanced excitability is widely spread through key brain areas. **(a, d, g)** Representative traces from neurons from three different brain regions-CA3 (top), BLA (middle) and mPFC (bottom). Recordings from F1-neurons of progenies of F0-Pseudo (cyan), F0-Trained (orange), and F0-Naïve parents (black) show effect of various training paradigms (or controls) over the amplitudes of the medium and late AHPs, summarized in **(b, c)** for CA3; **(e, f)** for BLA and **(h, i)** for mPFC. Data are presented as mean ± SEM. *, P<0.05; **, P<0.01; ***, p<0.001; n.s., non-significant.

**Supplementary Figure 5.**
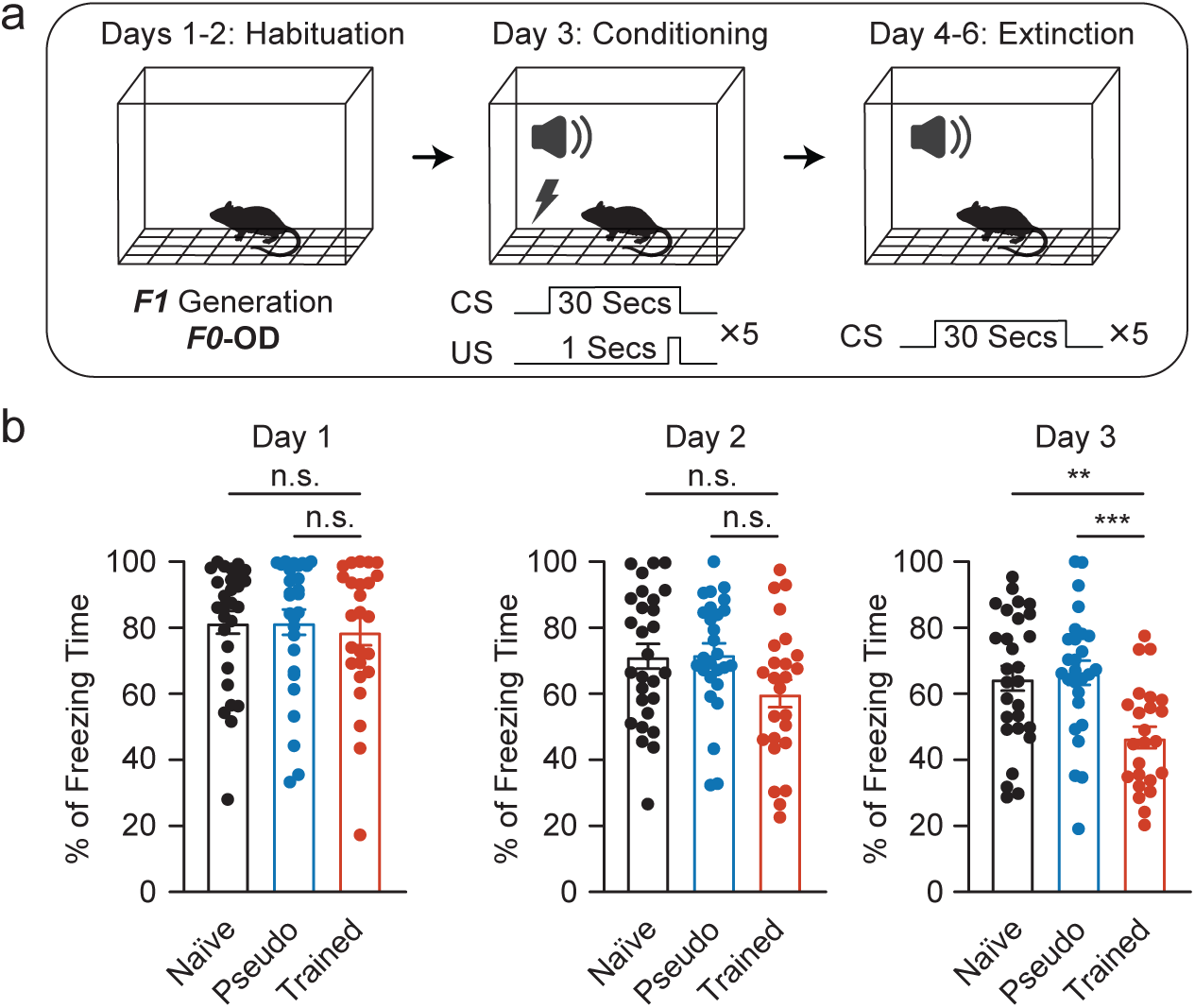
Enhanced performance of F1 progenies from OD-trained F0 rats in fear conditioning. **(a)** Illustration of the behavioral assay of fear conditioning and extinction. Freezing time (%) was measured for ten minutes during three consecutive days (see **methods**). **(b)** Offspring of trained rats show enhanced extinction of contextual fear conditioning (orange), in which rats learned that the context no longer predicts a following shock. Data are presented as mean ± SEM. *, P<0.05; **, P<0.01; ***, p<0.001; n.s., non-significant.

**Supplementary Figure 6.**
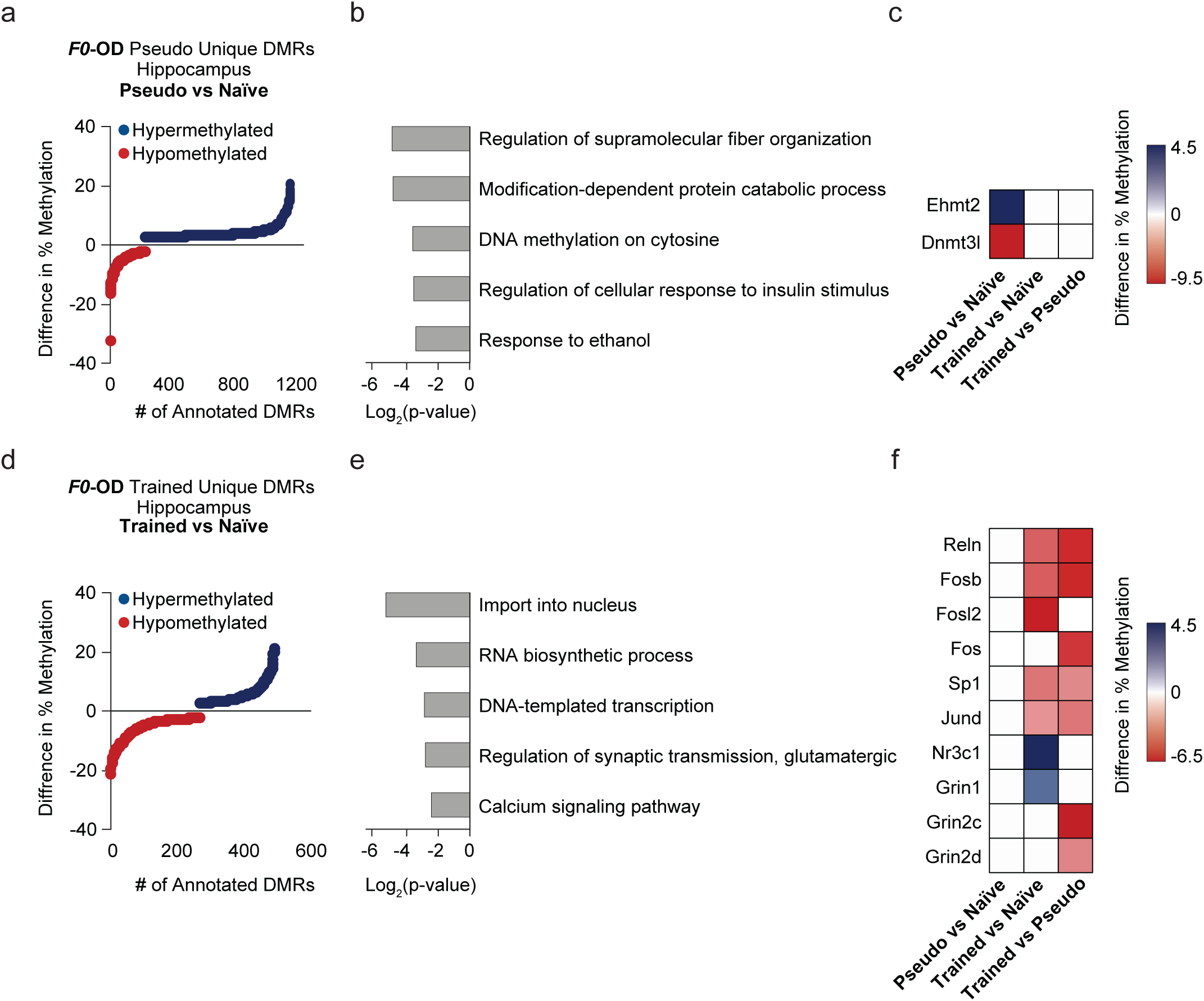
F0-Pseudo and F0-Trained unique DMRs gene ontology and genes. **(a, d)** Unique F0-pseudo or F0-Trained DMRs in rats. Each dot represents percentage of change in DMR (Hypermethylation in blue and hypomethylation in red) levels from the pair-wise comparison to F0-Naïve group. **(b, e)** Gene ontology (GO) analysis of unique F0-Pseudo or F0-Trained DMRs (via Metascape); values are presented in log_2_(*p-*value) scale. **(c, f)** Heatmap showing representative genes annotated to DMRs extracted from unique F0-Pseudo or F0-Trained DMRs GO analysis. Rows represent the percentage change in DMR levels from pairwise comparisons: Pseudo *vs* Naïve, Trained *vs* Naïve, and Trained *vs* Pseudo. Percent change in methylation values are color-coded and scaled: blue indicates hypermethylation, and red indicates hypomethylation.

**Supplementary Figure 7.**
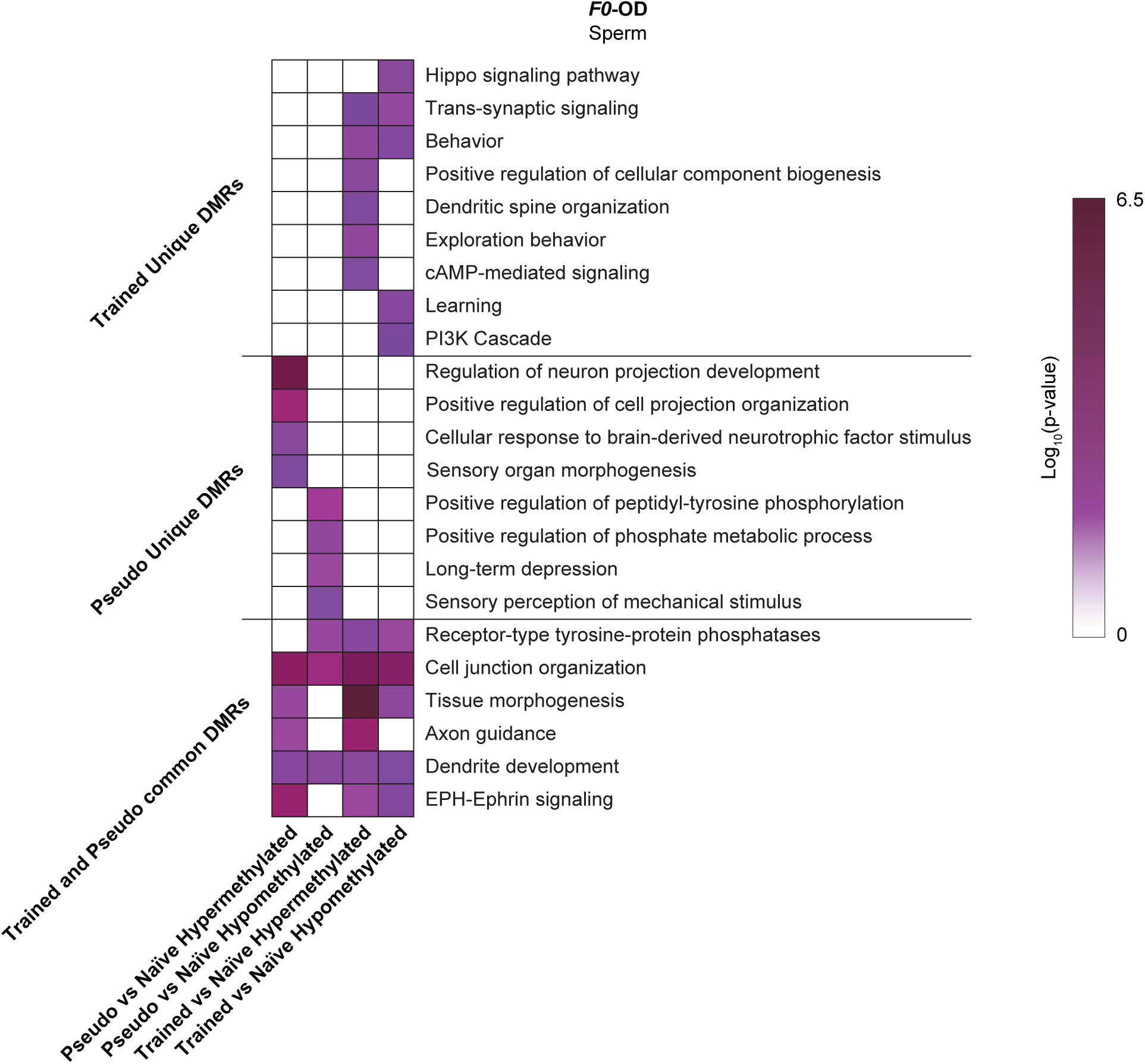
Gene ontology of Trained and Pseudo unique DMRs. **(a)** Heatmap displaying gene ontology (GO) enrichment of annotated DMRs from F0 sperm samples. Each row represents categories from Hypomethylated and Hypermethylated DMRs in pairwise comparisons of Pseudo vs Naïve and Trained vs Naïve. Values are presented on a log_10_(P value) color scale.

**Supplementary Table 1.** Overlap of Hypo- and Hyper-DMR–Annotated Genes in F0 and F1 Hippocampi (N = Naïve, P = Pseudo, T = Trained)

|  | lncRNA (Ensembl name) | Trained vs. Naive | Pseudo vs. Naive | Predicted genes |
| --- | --- | --- | --- | --- |
| Unique to trained | ENSMUST00000177421 | Hyper-DMR | - | Gsk3a,<br>Gsk3b,<br>Kcnq3,<br>Kcnq4,<br>Kcnq5, |
|  |  |  |  | Gnaq,<br>Plcb2,<br>Chrm5 |
|  | ENSMUST00000222673 | **Hypo-DMR** | - | Gsk3a,<br>Kcnq4,<br>Kcnq5,<br>Gnas,<br>Chrm4,<br>Chrm5 |
|  | ENSMUST00000183867 | **Hypo-DMR** | - | Gsk3b,<br>Gnaq,<br>Akap5 |
|  | ENSMUST00000212635 | **Hyper-DMR** | - | Calm1,<br>Plcl2 |
|  | ENSMUST00000210265 | **Hypo-DMR** | - | Plcb4 |
| Common | ENSMUST00000183867 | **Hypo-DMR** | **Hyper-DMR** | Gsk3b |
|  | ENSMUST00000133337 | **Hyper-DMR** | **Hyper-DMR** | Plcb2 |
| Unique to<br>pseudo | ENSMUST00000181534 | - | **Hyper-DMR** | Kcnq4,<br>Chrm4 |
|  | ENSMUST00000162770 | - | **Hypo-DMR** | Gnaq,<br>Chrm4,<br>Chrm5 |
|  | ENSMUST00000207592 | - | **Hyper-DMR** | Plcb4 |

